# Structural perturbation of chromatin domains with multiple developmental regulators can severely impact gene regulation and development

**DOI:** 10.1101/2024.08.03.606480

**Authors:** Shreeta Chakraborty, Nina Wenzlitschke, Matthew J. Anderson, Ariel Eraso, Manon Baudic, Joyce J. Thompson, Alicia A. Evans, Lilly M. Shatford‑Adams, Raj Chari, Parirokh Awasthi, Ryan K. Dale, Mark Lewandoski, Timothy J. Petros, Pedro P. Rocha

## Abstract

Chromatin domain boundaries delimited by CTCF motifs can restrict the range of enhancer action. However, disruption of domain structure often results in mild gene dysregulation and thus predicting the impact of boundary rearrangements on animal development remains challenging. Here, we tested whether structural perturbation of a chromatin domain with multiple developmental regulators can result in more acute gene dysregulation and severe developmental phenotypes. We targeted clusters of CTCF motifs in a domain of the mouse genome containing three FGF ligand genes—*Fgf3, Fgf4*, and *Fgf15*—that regulate several developmental processes. Deletion of the 23.9kb cluster that defines the centromeric boundary of this domain resulted in ectopic interactions of the FGF genes with enhancers located across the deleted boundary that are active in the developing brain. This caused strong induction of FGF expression and perinatal lethality with encephalocele and orofacial cleft phenotypes. Heterozygous boundary deletion was sufficient to cause these fully penetrant phenotypes, and strikingly, loss of a single CTCF motif within the cluster also recapitulated ectopic FGF expression and caused encephalocele. However, such phenotypic sensitivity to perturbation of domain structure did not extend to all CTCF clusters of this domain, nor to all developmental processes controlled by these three FGF genes—for example, the ability to undergo lineage specification in the blastocyst and pre-implantation development were not affected. By tracing the impact of different chromosomal rearrangements throughout mouse development, we start to uncover the determinants of phenotypic robustness and sensitivity to perturbation of chromatin boundaries. Our data show how small sequence variants at certain domain boundaries can have a surprisingly outsized effect and must be considered as potential sources of gene dysregulation during development and disease.

## INTRODUCTION

Successful embryonic development requires dynamic spatial and temporal expression of genes in a cell type specific manner. This is particularly important for genes that encode developmental regulators of processes such as embryonic patterning or cell specification. Gene expression is regulated by enhancers that can act across large distances from their target genes (de Laat and Duboule, 2013; Furlong and Levine, 2018; Long et al., 2016; Schoenfelder and Fraser, 2019). The spatial range of enhancer action has been proposed to be delimited by topologically associated domains (TADs), which are characterized by higher levels of intra-domain contact frequency compared to neighboring chromatin regions (Beagan and Phillips-Cremins, 2020; Dixon et al., 2012; Mirny and Dekker, 2022; Misteli, 2020; Nora et al., 2012; Rowley and Corces, 2018). TADs can prevent enhancer off-target gene activation by reducing the frequency of contacts across their boundaries. In vertebrates, these chromatin domains are mostly formed through the process of loop extrusion by the cohesin complex, where domain boundaries are frequently delimited by CTCF, which retains extruding cohesin complexes in an orientation-dependent manner determined by the sequence of its DNA binding motif (de Wit et al., 2015; Fudenberg et al., 2016; Nora et al., 2017; Rao et al., 2017; Schwarzer et al., 2017; Zhang et al., 2020). The boundaries of these chromatin domains are often composed of clusters of multiple CTCF binding motifs in both orientations, which can increase the chances of cohesin retention and thus generate better insulation (Anania et al., 2022; Chang et al., 2023).

The loop extrusion model predicts that perturbation of the extrusion machinery, or sequence variants that disrupt CTCF binding, can result in loss of insulation, ectopic enhancer-promoter interactions, and off-target gene activation with potential pathogenic outcomes. Indeed, such effects have been observed in multiple studies (Baudic et al., 2024; Cova et al., 2023; Dowen et al., 2014; Ealo et al., 2023; Engel et al., 2008; Flavahan et al., 2016; Franke et al., 2016; Hanssen et al., 2017; Juan et al., 2022; Kraft et al., 2019; Lupianez et al., 2015; Narendra et al., 2015; Rajderkar et al., 2023; van Bemmel et al., 2019; Wu et al., 2021). However, disruption of chromatin domains does not always impact gene regulation or affect physiological processes. For example, in vitro, rapid degradation of cohesin or CTCF leads to the dissolution of TADs but the impact on transcription is minimal (Nora et al., 2017; Rao et al., 2017). Disruption of some chromatin domains has a negligible effect on gene regulation and animal development (Amandio et al., 2021; Despang et al., 2019; Ghavi-Helm et al., 2019; Paliou et al., 2019; Rajderkar et al., 2023; Rodriguez-Carballo et al., 2020; Williamson et al., 2019). Our own previous work showed that some *Sox2* enhancers can bypass strong ectopic CTCF-mediated boundaries, which contributes to phenotypic robustness (Chakraborty et al., 2023). In sum, we do not yet fully understand the physiological significance of CTCF-mediated boundaries in orchestration of gene regulation and ensuring successful embryonic development.

Here, we asked whether chromatin domains with multiple developmental regulators require better insulation and therefore may be more susceptible to cause severe developmental phenotypes upon perturbation of their boundaries. To test this, we targeted a chromatin domain that harbors three FGF ligand genes—*Fgf3, Fgf4*, and *Fgf15*—separated from each other by clusters of CTCF motifs. Like other members of this large family of signaling ligands (Brewer et al., 2016), these genes have distinct expression patterns and developmental functions. We show that disruption of CTCF-mediated insulation can result in remarkable developmental phenotypes with full penetrance even in heterozygosity. In fact, loss of CTCF binding at a single motif within a large cluster was enough to severely dysregulate gene expression and impair mouse development. Importantly, we did not observe such a strong phenotypic impact from perturbing other CTCF clusters of this chromatin domain, and many developmental processes controlled by these FGF ligands were not affected in any of our mutants. In summary, by tracing the impact of different structural rearrangements throughout development, we start to uncover determinants of phenotypic robustness and sensitivity to perturbation of chromatin boundaries. Our work also emphasizes how development and gene regulation can be completely intolerant of small sequence variants at specific chromatin boundaries.

## RESULTS

### Deletion of CTCF clusters in a chromatin domain with three FGF genes disrupts mouse development

*Fgf3, Fgf4* and *Fgf15* reside within a 63kb region of the mouse chromosome 7 and are separated from each other by clusters of CTCF motifs (**Fig. 1A**). Upstream of *Fgf3*, there is a 23.9kb cluster with four CTCF motifs that we named C1-C4. The 6kb C5-C6 cluster is located between *Fgf3* and *Fgf4*, and immediately upstream of *Fgf15*, the C7-C10 cluster occupies 8.5kb. The C11-C13 cluster separates *Fgf15* from *Lto1* and *Ccnd1*. Each cluster has CTCF motifs in both forward and reverse orientations and ChIP-seq in mouse embryonic stem (ES) cells confirmed co-occupancy of the cohesin subunit RAD21 at all CTCF sites except C8, C11 and C13. Capture-HiC (CHi-C) in ES cells showed that the three FGF genes reside within a highly interacting domain delimited by C1-C4 on its centromeric end. (**Fig. 1A**). The C5-C6 cluster overlaps a weak boundary between *Fgf3* and *Fgf4*, and C7-C10 also does not completely insulate *Fgf15* from *Fgf3* and *Fgf4* as some interactions can be observed across this cluster. In contrast, *Fgf15* is well insulated from downstream genes including *Lto1* and *Ccnd1*, in a boundary delimited by the C11-C13 cluster that marks the telomeric end of this domain.

**Figure 1.**
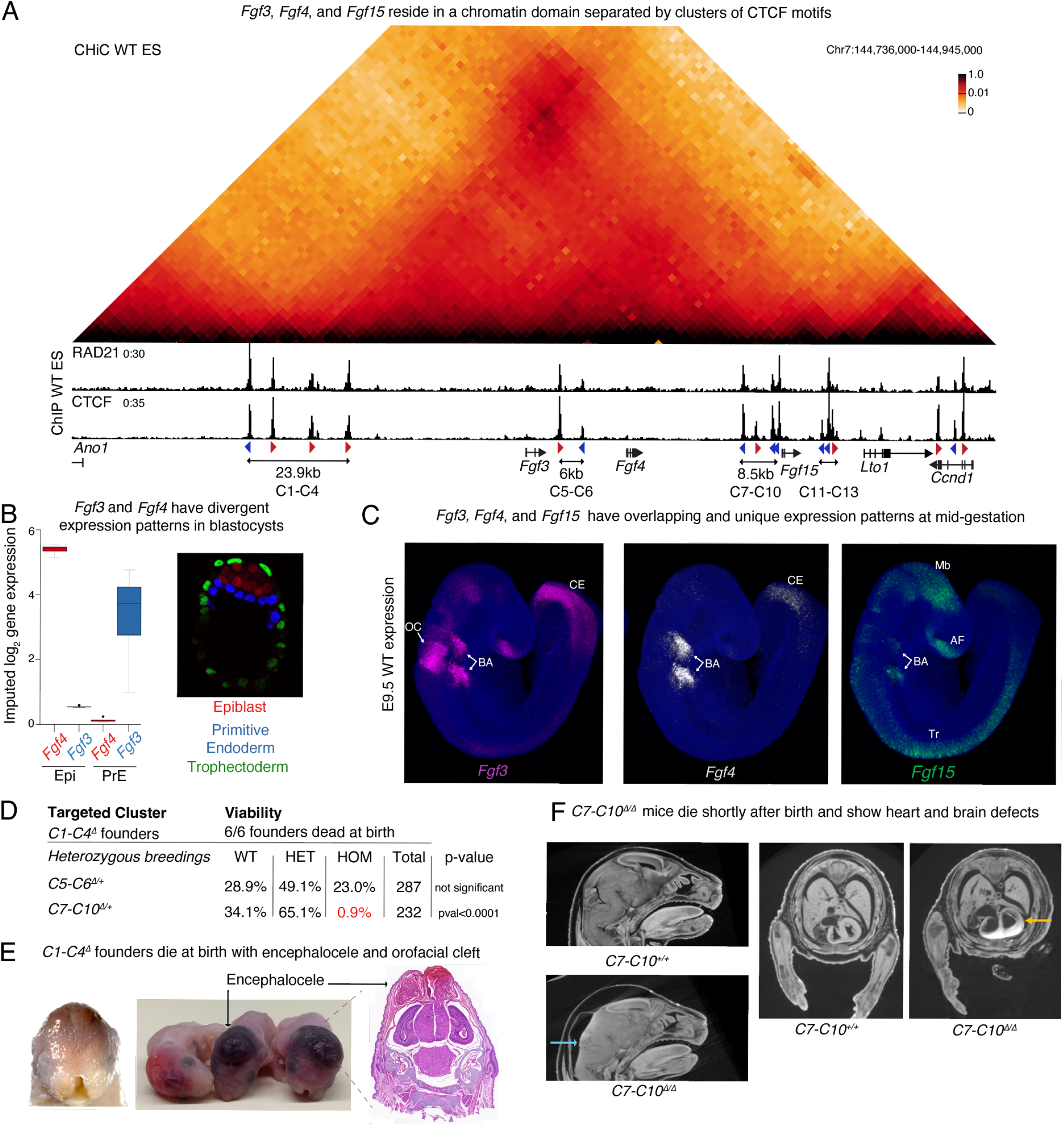
Deletion of CTCF clusters in a chromatin domain with three FGF genes causes embryonic lethality. **A** CHi-C 1D interaction frequency heatmap at 2kb resolution (top) and ChlP-seq for RAD21 and CTCF in wild-type mES cells (bottom). CTCF motif orientation (red and blue arrowheads) is shown for significant CTCF motifs (pval <0.05) as detected by FIMO. **B** Expression of *Fgf3* and *Fgf4* in single cells labelled as Epi or PrE in E4.5 blastocysts (left) as quantified in endoderm-explorer.com. Representative image of the three cell-types in mouse blastocysts at E4.5 (right) labelled by antibodies against specific markers: Trophectoderm-CDX2; PrE-GATA6; Epi-NANOG. **C** Expression of the 3 FGF genes in E9.5 mouse embryos as labeled by HCR (n=3/3). AF-Anterior forebrain, BA-Branchial Arches, CE-Caudal end, Mb-Midbrain, OC-Otic cup, Tr-Trunk **D** Viability of the three mouse lines with CTCF cluster deletions. All founders with at least one copy of the *C1-C4*^*Δ*^died perinatally. Viability of pups generated from crosses of heterozygotes of the *C5-C6*^*Δ*^ and *C7-C10*^*Δ*^ mouse lines was assessed at weaning. No deviation from expected mendelian ratio was found for *C5-C6*^*Δ/Δ*^ homozygotes. Only 2 *C7-C10*^*Δ/Δ*^ homozygous pups were recovered alive at weaning out of 58 expected. **E** Representative image of *C1-C4*^Δ^founder pups showing encephalocele and orofacial cleft (left) (n=6/6). H&E staining on a paraffin frontal section of a *C1-C4*^*Δ*^ pup. **F** Brain and heart defects seen in *C7-C10*^*Δ/Δ*^ homozygotes using Micro-CT scans. Blue arrow points to fused ventricles with edema and yellow arrow points to enlarged heart ventricles (n=6/7).

These FGF genes exhibit diverse tissue-specific expression patterns and functions. *Fgf4* is essential for cell fate specification in the blastocyst, and successful post-implantation development (Feldman et al., 1995; Kang et al., 2013). At embryonic day 4.5 (E4.5), publicly available single-cell RNA-seq data (Nowotschin et al., 2019) show that *Fgf3* is expressed in cells classified as primitive endoderm (PrE) of blastocysts, while *Fgf4* is specifically found in epiblast (Epi) cells (**Fig. 1B**). At mid-gestation (E9.5), in situ hybridization chain reaction (HCR) analysis demonstrated that the FGF genes have not only tissue-specific expression patterns but also overlapping ones. For example, the branchial arches expressed all three FGF ligands but higher levels of *Fgf3* and *Fgf4*. Combined loss of *Fgf3* and *Fgf4* in the branchial arches causes lethality (Zelarayan et al., 2022). *Fgf3* is the only gene out of the three that is highly expressed in the otic cup and contributes to patterning of the inner ear (Mansour et al., 1993) (**Fig. 1C**). *Fgf15* is strongly expressed in the midbrain and anterior forebrain, where it is required for normal development, as well as the branchial arches where it acts on the cardiac neural crest (Borello et al., 2008; Vincentz et al., 2005). In the trunk, both *Fgf3* and *Fgf4* are highly expressed towards the caudal end, while *Fgf15* is found more proximally in a non-overlapping domain.

Since the three FGF genes achieve different patterns of tissue-specific expression despite their genomic proximity, we hypothesized that the CTCF clusters separating them are required to insulate each other’s regulatory elements. To test this, we generated mouse lines where we deleted the CTCF clusters that are found between FGF genes (C5-C6 and C7-10). Furthermore, to test if perturbing the structure of a domain with multiple developmental regulators would cause severe phenotypes, we decided to target one of the clusters that marks a boundary of this domain and chose C1-C4 at the centromeric end. These three mouse lines were generated by injecting zygotes with Cas9 and gRNAs fully surrounding each cluster to ensure complete loss of cohesin retention and disruption of insulation (**Fig. S1A**). All founder mice with an allele where the C1-C4 cluster was deleted (*C1-C4*^*Δ*^) were found dead at birth, despite repeated injection attempts. In contrast, we successfully established mouse lines with deletions of the C5-C6 and C7-C10 clusters (*C5-C6*^*Δ*^ and *C7-C10*^*Δ*^, respectively). Crosses between F1 *C5-C6*^*Δ/+*^ heterozygous animals resulted in recovery of homozygotes at the expected Mendelian ratio (**Fig. 1D**). Strikingly, almost all *C7-C10*^*Δ/Δ*^ homozygotes died perinatally. To understand how individual CTCF cluster deletions affected viability and development, we investigated the cause of lethality. *C1-C4*^*Δ*^ founder mice exhibited craniofacial abnormalities with orofacial clefts (**Fig. 1E**). H&E staining of frontal sections of the head demonstrated the presence of encephalocele, a neural tube closure defect where neural tissues protrude outside the skull and are covered with an epithelial membrane. Postnatal lethality of *C7-C10*^*Δ/Δ*^ homozygotes could not be attributed to specific developmental abnormalities but Micro-CT scan of E18.5 fetuses revealed that homozygous mutants had a range of phenotypes including occluded and reduced brain vesicles as well as enlarged heart ventricles (**Fig. 1F**). In sum, unlike disruption of CTCF-mediated chromatin boundaries at other loci (Chakraborty et al., 2023; Despang et al., 2019; Rajderkar et al., 2023; Williamson et al., 2019), deletion of CTCF clusters in a chromatin domain with multiple developmental regulators severely impacted animal development.

### Cell type-specific expression of FGF genes in blastocysts does not rely on CTCF-mediated insulation

Although *Fgf3* and *Fgf4* are close genomic neighbors, they are divergently expressed in mouse blastocysts. *Fgf3* is found exclusively in the PrE, while *Fgf4* is an Epi marker (**Fig. 1B**). The chromatin environment of these genes also display contrasting patterns. ES cells, which are an Epi in vitro model, are enriched with H3K27ac at *Fgf4*—including on the 3’ UTR where sequences that can drive Epi expression have been identified (Fraidenraich et al., 1998)—while *Fgf3* is decorated with the repressive mark H3K27me3. In stark contrast, in vitro models of the PrE (XEN cells) have the exact opposite pattern of chromatin marks at these two genes (**Fig S2A**). Because tight regulation of FGF4 levels is essential for post-implantation development, we expected that deletion of the CTCF clusters on either side of *Fgf4—* in the *C5-C6*^*Δ*^ and *C7-C10*^*Δ*^ lines—would affect expression of the FGF ligands and perturb implantation. However, all mutant mice successfully implanted (**Fig. 1D**). We therefore decided to focus on this early developmental stage to better understand the mechanisms underlying phenotypic resilience to perturbation of CTCF boundaries in some developmental contexts. We first assessed if deletion of the C5-C6 and C7-C10 clusters causes ectopic activation of the FGF genes located across each deleted boundary using RNA-seq in single blastocysts. Surprisingly, we did not detect any of the three FGF ligand genes in either *C5-C6*^*Δ/Δ*^ or *C7-C10*^*Δ/Δ*^ homozygotes to be differentially expressed (false discovery rate (FDR) <0.1 and log_2_FC>1) (**Fig. 2A**). This suggests that the enhancers that control expression of *Fgf3* and *Fgf4* in blastocysts, do not rely on the CTCF clusters to achieve specificity of target gene induction.

**Figure 2.**
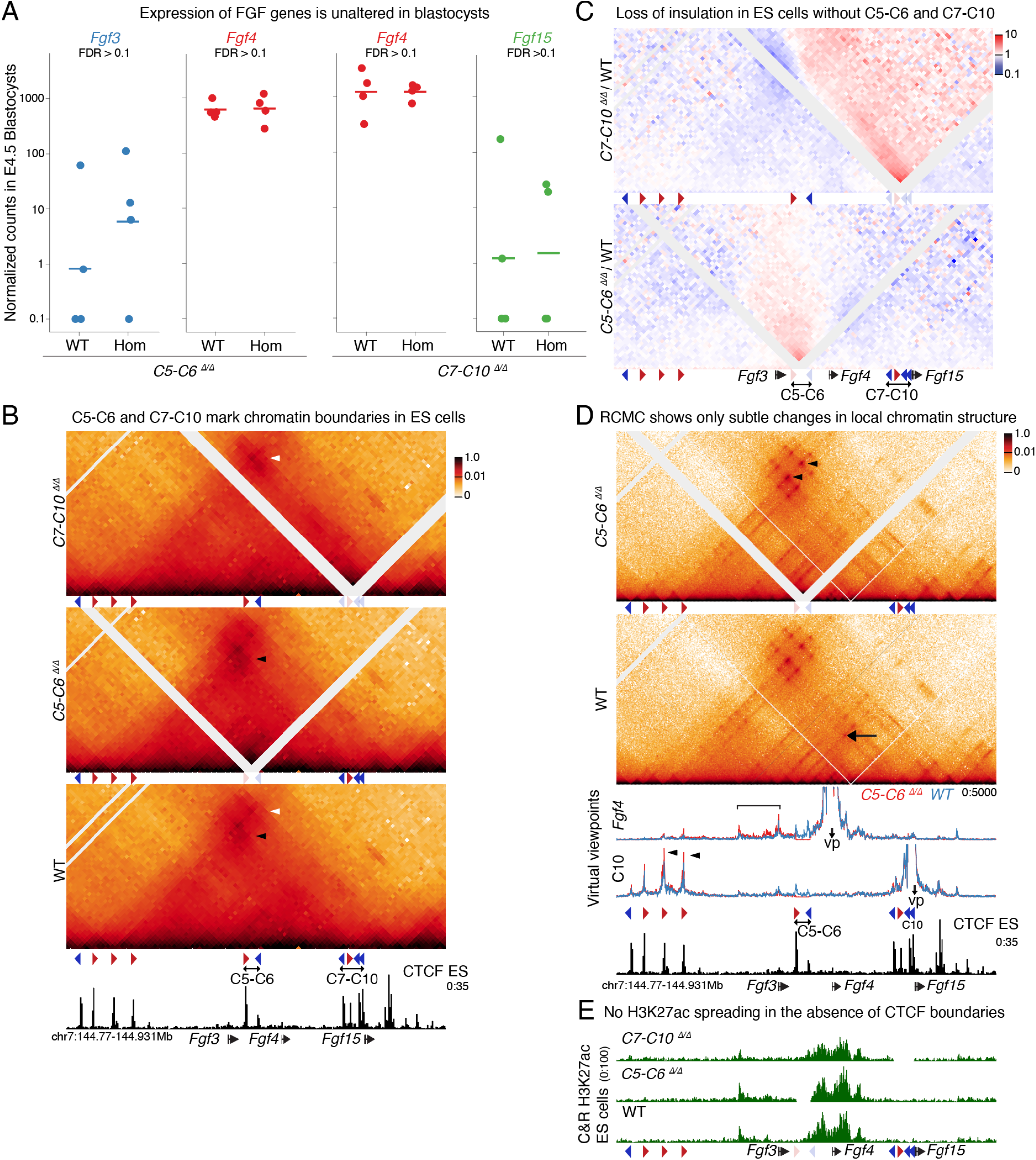
Tissue-specific expression of *Fgf3* and *Fgf4* in blastocysts does not require CTCF-mediated insulation. **A** Expression of FGF genes in single blastocysts as measured by RNA-seq in *C5-C6*^*Δ/Δ*^ and *C7-C10*^*Δ/Δ*^ homozygous compared to WT littermates. Each circle represents a blastocyst and the bar represents the median DESeq2-normalized expression value of each genotype. No gene was identified as significantly differently expressed (FDR <0.1, log_2_fc>0.1). **B** CHiC 1D interaction frequency heatmap in *C5-C6*^*Δ/Δ*^ and *C7-C10*^*Δ/Δ*^ homozygous mES cells, compared to WT at 2kb resolution. Arrowheads represent increased focal interactions between the CTCF clusters that surround the deleted clusters. White diagonal lines reflect regions without mapped reads caused by deletions or failure to map to repetitive regions. **C** Differential CHiC interaction frequency heatmap. Red signal represents interactions that occur at higher frequency in mutant cell lines compared to control and blue shows interactions of lower frequency. **D** RCMC 1D interaction frequency heatmap in *C5-C6*^*Δ/Δ*^ homozygous ES cells compared to WT at 400bp resolution. Below heatmaps, data are shown as virtual viewpoints at 50bp resolution Viewpoints were centered on the *Fgf4* promoter and the C10 CTCF motif Black arrow shows interaction between C5 and the C7-C10 cluster. Black arrowheads highlight increased interactions between C1-C4 and C7-C10 in *C5-C6*^*Δ/Δ*^ cell. Bracket highlights increased interactions from the *Fgf4* viewpoint with regions across the deleted boundary. White diagonal lines reflect regions without mapped reads caused by deletions or failure to map to repetitive regions. **E** CUT&RUN H3K27ac data in mES cells of three different genotypes.

Since we did not observe changes in gene expression in these three genes, we first verified whether the C5-C6 and C7-C10 clusters can function as CTCF loop anchors capable of insulating interactions. This was especially relevant for the C5-C6 cluster, as wild-type ES cells do not display strong insulation at this region (**Fig. 1A**). Since the low cell numbers of blastocysts impair high-resolution confirmation capture techniques, we used ES cells as proxies for the Epi state. We derived ES cell lines from *C5-C6*^*Δ/Δ*^ and *C7-C10*^*Δ/Δ*^ homozygous blastocysts in parallel with wild-type littermates and performed CHi-C (**Fig. 2B**). Loss of CTCF motifs in *C5-C6*^*Δ/Δ*^ cells led to increased focal interactions between the upstream cluster C1-C4 with C7-C10 (black arrowhead in **Fig. 2B**). Similarly, deletion of motifs in *C7-C10*^*Δ/Δ*^ enhanced contacts between C1-C4 and the CTCF cluster downstream of *Fgf15* (white arrowhead in **Fig. 2B**). Differential interaction frequency heatmaps highlight the decrease in insulation and increase in contacts across the deleted clusters (**Fig. 2C**). In a complementary approach, we differentiated *C5-C6*^*Δ/Δ*^ ES cells and matched wild-type controls into XEN cells that express *Fgf3*. XEN cells also lost insulation between chromatin domains and increased contacts between neighboring CTCF clusters (**Fig. S2B; S2C**). Together, these data demonstrate that both C5-C6 and C7-C10 CTCF clusters function as loop anchors that can insulate interactions between chromatin domains.

To identify potential ectopic enhancer–promoter interactions caused by loss of CTCF-mediated insulation we generated Region Capture Micro-C (RCMC) contact maps that have higher resolution than CHi-C and allow better identification of regulatory interactions (**Fig. 2D**) (Goel et al., 2023). RCMC again confirmed that C5-C6 functions as a chromatin domain boundary. As observed with CHi-C, *C5-C6*^*Δ/Δ*^ ES cells had slightly elevated contact frequency between CTCF motifs of the C1-C4 and the C7-C10 cluster (black arrowheads **Fig. 2D**). Furthermore, these high resolution data—using 50bp bins—demonstrate that the C5 motif serves as an anchor in loops with motifs in the C7-C10 cluster (black arrow in WT heatmap **Fig. 2D**). To characterize changes in enhancer activity we complemented RCMC with H3K27ac Cut&Run (**Fig. 2E**). Despite the loss of insulation, contacts between the active *Fgf4* promoter and enhancers across this chromatin domain were only slightly increased in *C5-C6*^*Δ/Δ*^ cells (bracket in *Fgf4* virtual viewpoints in **Fig. 2D; S2D**) and no ectopic enhancer-promoter contacts were found. Furthermore, the pattern of H3K27ac in *C5-C6*^*Δ/Δ*^ and *C7-C10*^*Δ/Δ*^ cells showed no spreading of active chromatin marks and mostly accumulated proximal to *Fgf4*. (**Fig. 2E**). In summary, these data suggest that blastocyst expression of *Fgf3* and *Fgf4* is mostly driven by extremely target-specific proximal enhancers independently of CTCF, which may contribute to the resilience of blastocysts to perturbation of chromatin structure during mammalian implantation.

### *C7-C10*^*Δ/Δ*^ phenotypes likely arise from perturbation of chromatin structure and deletion of enhancers

In contrast to the phenotypic robustness of blastocysts to perturbation of chromatin structure, loss of CTCF binding in the C1-C4 and C7-C10 clusters was remarkably impactful later in development. We first asked whether cluster deletions affected only CTCF binding or if phenotypes could be caused by loss of other types of regulatory elements. We first tried to understand the cause of the heart and brain defects of *C7-C10*^*Δ/Δ*^ homozygotes (**Fig. 1F**). For this, we assessed expression of *Fgf3, Fgf4* and *Fgf15* at E9.5 using HCR and saw that expression of *Fgf15* in the brain, and of all three FGF genes in the branchial arches was severely reduced (**Fig. S3A**). Due to the physical proximity of the *C7-C10*^*Δ*^ deletion to *Fgf15*, we wondered if putative enhancer elements could have inadvertently been deleted in this allele. Analysis of publicly available ChIP-seq and ATAC-seq in wild-type developing brain revealed enrichment of H3K27ac and accessible chromatin regions that partially overlapped with the C7-C10 deletion (**Fig. S3B**). Furthermore, this region contains sequences that recapitulate some of the expression patterns of *Fgf15* (Saitsu et al., 2006). Along with the loss of *Fgf3, Fgf4*, and *Fgf15* expression in the brain and branchial arches, this strongly suggests that the phenotypes seen in *C7-C10*^*Δ/Δ*^ homozygotes may not solely be caused by loss of CTCF binding and could also be due to deletion of enhancers. Importantly, *Fgf15* expression was not disrupted in all tissues. In fact, *C7-C10*^*Δ/Δ*^ homozygotes showed ectopic expression of *Fgf15* in the apical ectodermal ridge of forelimbs (**Fig. S3C**). This confirmed that the core promoter elements of *C7-C10*^*Δ/Δ*^ animals were not disrupted. Unlike *C7-C10*^*Δ/Δ*^ pups that die at birth, animals without *Fgf15* survive until weaning (Borello et al., 2008; Vincentz et al., 2005). Since the defects we report here are more severe than those described for the *Fgf15* knock out, it is likely that the developmental defects of *C7-C10*^*Δ/Δ*^ mice are caused by a combination of insulation loss and deletion of enhancers within the C7-C10 cluster. Because distinguishing between these two effects would be very challenging, we switched our focus to the CTCF cluster deletion at the centromeric end of this domain.

### Loss of CTCF binding at the C1-C4 cluster causes perinatal heterozygous lethality

On the centromeric end of the domain, the heterozygous lethality of pups with the *C1-C4*^*Δ*^ allele, created an obstacle to study the function of this cluster in animal development. To overcome this and assess whether the phenotypes observed in mice with the *C1-C4*^*Δ*^ allele resulted from loss of CTCF binding, we attempted a genetic rescue of the embryonic defects (**Fig. 3A**). We used the same gRNAs that generated the *C1-C4*^*Δ*^ allele (**Fig. S1B**) along with a 672bp repair template containing the four CTCF motifs found in the C1-C4 boundary, flanked by *loxP* sites—an allele we named *C1-C4*^*flx*^. Strikingly, despite deleting 23.9kb, reintroduction of these four CTCF motifs was sufficient to rescue lethality, and *C1-C4*^*flx/flx*^ homozygotes were both viable and fertile (**Fig. 3B**). We then bred *C1-C4*^*flx/+*^ heterozygous mice with homozygous *E2A-Cre*^*t/t*^ animals that ubiquitously express Cre recombinase to generate embryos with heterozygous loss of the C1-C4 cluster (*C1-C4*^*flx/+*^; *E2A-Cre*^*t*^) (Lakso et al., 1996). These embryos recapitulated the encephalocele and orofacial cleft defects in a fully penetrant manner and died perinatally (**Fig. 3C; 3D; S3D**). The rescue of the severe developmental phenotypes in mice with the floxed CTCF cassette, and the recapitulation of these phenotypes upon Cre recombination provides direct evidence that the defects observed in *C1-C4*^*Δ*^ mice are caused by loss of CTCF-binding and not by deletion of other regulatory elements. It also highlights the sensitivity of mouse development to loss of CTCF binding at this domain boundary, since heterozygous deletion of the CTCF cluster is sufficient to cause such strong developmental phenotypes and fully penetrant perinatal lethality.

**Figure 3.**
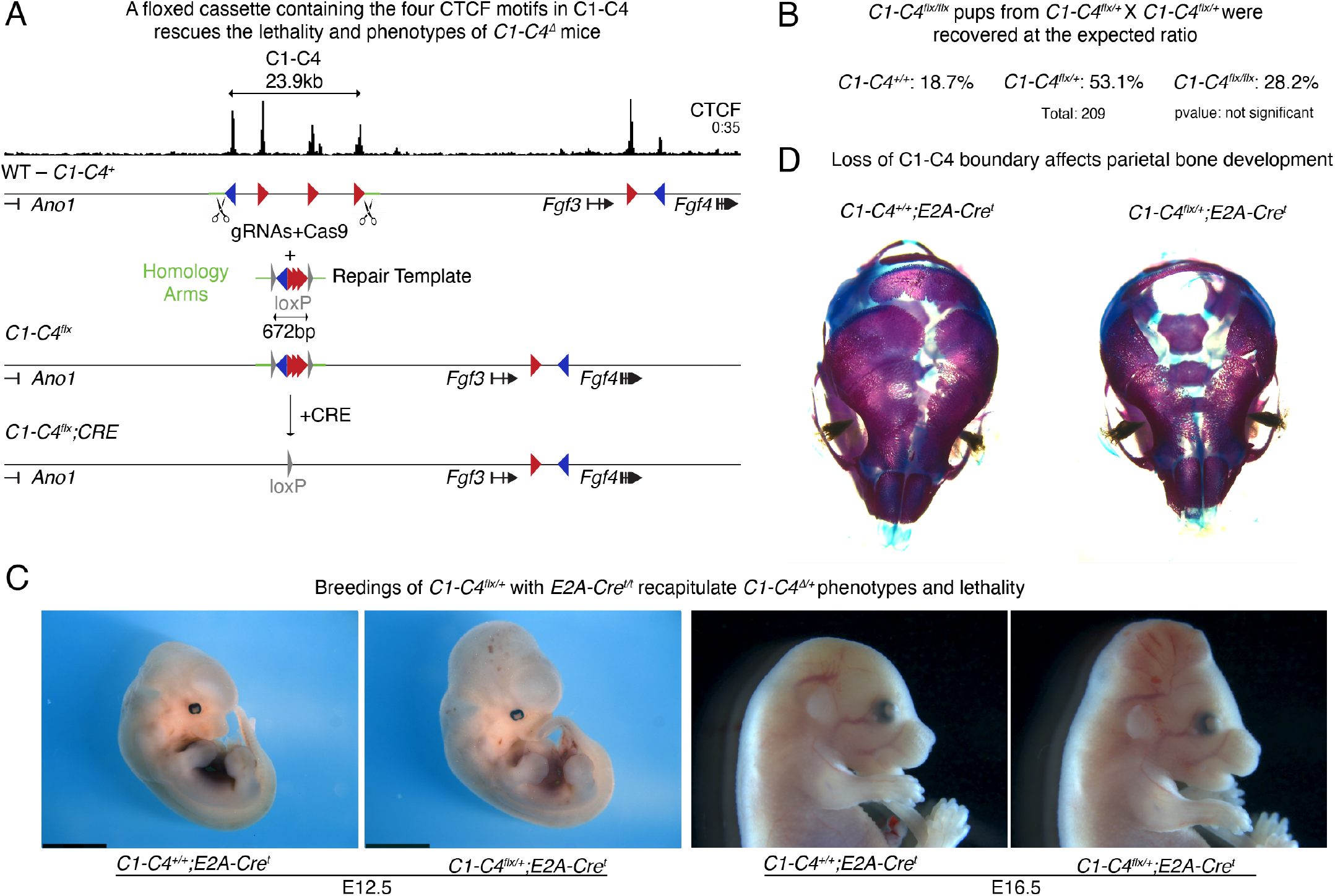
Heterozygous deletion of the C1-C4 cluster causes lethality because of loss of CTCF binding. **A** Scheme Of the Strategy used to generate the *C1-C4*^*flx*^ allele and result from Cre-mediated recombination. Green lines represent regions of homology present in the repair template. Repair template contains also aproximatelly 120bp from each of the four CTCF motifs of the C1-C4 boundary flanked by loxP sites (gray triangles). Upon Cre-mediated recombination of the cassette in mice with the *C1-C4*^*flx*^ cassette, the four CTCF motifs are lost and a single loxP motif is left behind. **B** Pups resulting from crosses of *C1-C4*^*flx/\**^ heterozygotes recover all alleles at expected mendelian ratio. This confirms that the floxed C1-C4 cassette fully rescues the phenotypes of *C1-C4*^*Δ*^ mice. **C** Embryos and fetuses resulting from the breeding of heterozygous *C1-C4*^*flx/+*^ with *E2A-Cre*^*t/t*^ homozygotes fully recapitulate the phenotypes seen in *C1-C4*^*Δ*^. At 12.5 the midbrain tissue appears expanded (left), while at 16.5 it is already outside of the skeletal tissue (n=6/6). **D** Skeletal preparations of the same crosses show a defect in parietal bone development (n=3/3).

Encephalocele is a severe birth defect found in one out of 10,500 human births (Mai et al., 2019), but its etiology is poorly understood in part because existing mouse models recapitulate the features that characterize the human disease with low penetrance (Rolo et al., 2019). To improve our understanding of this phenotype, we examined *C1-C4*^*flx/+*^; *E2A-Cre*^*t*^ heterozygous embryos at E12.5 and E16.5. This revealed an accumulation of fluid in mutant heads, over-expansion of neural tissue (**Fig. 3C**), and abnormal parietal bone development (**Fig. 3D**). It was unclear if the bone pathology was caused by secondary effects of brain herniation, ossification defects or anomaly in skull bone fusion. Because craniofacial phenotypes are often caused by defects in neural crest cells, we next investigated if defects in this cell type contributed to the phenotypes. To discern the origin of the phenotypes, we crossed the *C1-C4*^*flx/+*^ heterozygotes with either *Wnt1-Cre* or *Sox10-Cre* mice. The *Wnt1-Cre* line expresses Cre throughout the neural tube including neural crest cells, while the *Sox10-Cre* line is restricted to migratory neural crest cells (Jacques-Fricke et al., 2012). All double heterozygotes, *C1-C4*^*flx/+*^; *Wnt1-Cre*^*t*^ showed perinatal lethality and recapitulated both encephalocele and cleft defects (**Fig. S3E**). Conversely, none of these phenotypes were observed in *C1-C4*^*flx/+*^; *Sox10-Cre*^*t*^ progeny (**Fig. S3F**). These data strongly suggest that the craniofacial phenotypes do not originate from migratory neural crest cells.

### Disruption of the C1-C4 boundary leads to ectopic expression of FGF genes in the brain

To understand the molecular mechanisms that caused pronounced phenotypes when CTCF binding was lost at the C1-C4 cluster, we bred *C1-C4*^*flx*^ mice with *ROSA26-Cre*^*ERT2*^ mice. This line ubiquitously expresses the Cre recombinase with its activity controlled by tamoxifen treatment. *C1-C4*^*flx/flx*^ homozygous females with one copy of the *Cre*^*ERT2*^ allele were then mated with *C1-C4*^*flx/flx*^ homozygous males and tamoxifen was orally administered to the females at E6.5 (see breeding scheme in **Fig. S4A**). This generated embryos that carried the intact *C1-C4*^*flx*^ allele in both chromosomes (referred to as *C1-C4*^*flx/flx*^) as well as littermates where the C1*-*C4 rescue cassette was deleted in homozygosity (referred to as *C1-C4*^*flx/ flx*^; Cre). As the midbrain was visibly enlarged at E12.5 (**Fig. 3B**), we collected this brain region at E11.5 when defects are not yet obvious. Subsequently, we performed CHi-C on each genotype and reads were aligned to a custom mm10 genome where the 23.9kb C1-C4 cluster was replaced by the 672bp C1-C4^flx^ cassette. This confirmed that the *C1-C4*^*flx/ flx*^ rescue cassette can function as an anchor capable of establishing loops with both upstream and downstream CTCF clusters (white arrowheads in **Fig. 4A**). Littermates with Cre-induced C1-C4 deletion showed complete loss of upstream and downstream loops, with increased interactions from flanking CTCF sites on both ends in the absence of the C1-C4 anchor (black arrowheads in **Fig. 4A**). Importantly, the loss of insulation caused fusion of the domain with the three FGF genes with the upstream domain containing the *Ano1* gene.

**Figure 4.**
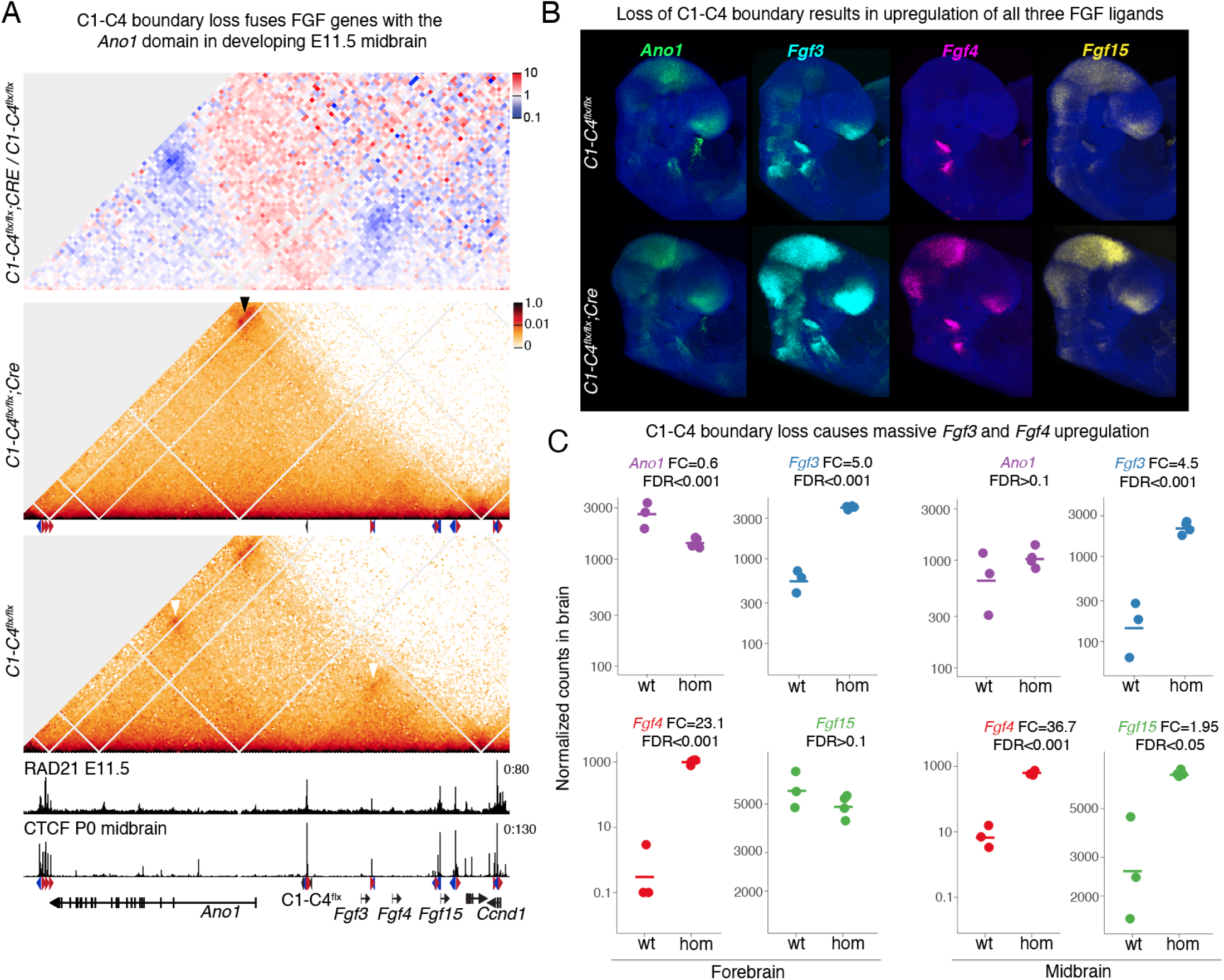
Disruption of the C1-C4 boundary leads to ectopic expression of FGF genes in the brain. **A** Differential CHiC interaction frequency heatmap at 4kb resolution between *C1-C4*^*flx/flx*^;CREand *C1-C4*^*flx/flx*^ microdissected E11.5 midbrains. Red signal represents interactions that occur at higher frequency in mutant cell lines compared to control and blue shows interactions of lower frequency (top). CHi-C 1D interaction frequency heatmap in *C1-C4*^*flx/flx*^;CRE and *C1-C4*^*flx/flx*^ microdissected E11.5 midbrains (bottom). All data, including RAD21 and CTCF CHiP-seq, were mapped to a custom mm10 genome where the the C1-C4 cluster was replaced by the *C1-C4*^*flx*^ rescue cassette. Arrowheads represent increased focal interactions between the CTCF clusters that surround the deleted clusters. **B** HCR of E9.5 embryos showing higher expression of FGF genes in the midbrain and anterior forebrain (n=4 for each genotype), **c** Expression of the three FGF genes and *Ano1* in micro-dissected midbrain and anterior forebrain at Ell .5 measured by RNA-seq. Each circle shows the expression of one embryo and bar represents the median value of each genotype. Fold change and FDR is shown above the plot.

Next, we examined the expression of genes flanking the C1-C4 cluster at E9.5 using HCR. This stage was chosen to facilitate imaging since the additional fluid accumulation in the heads of all E10.5 mutant embryos leaked during embryo processing for HCR which resulted in collapsed heads (data not shown). Loss of the C1-C4 rescue cassette strongly increased *Fgf3, Fgf4* and *Fgf15* expression in the midbrain and anterior forebrain (**Fig. 4B**). Interestingly, all three FGF genes in mutant brains recapitulated the expression pattern of *Ano1*, a gene that encodes a chloride ion channel located in the upstream domain. We then bred homozygous *C1-C4*^*flx/ flx*^ males with homozygous E2A-Cre females to generate embryos with heterozygous loss of the C1-C4 boundary and control littermates. Importantly, these heterozygous embryos also showed the striking upregulation of the FGF genes (**Fig. S4B**). To better quantify changes in expression of the FGF genes and match expression data with CHi-C, we used RNA-seq on micro-dissected E11.5 midbrains and anterior forebrains. As indicated by HCR, *Fgf4* exhibited the most dramatic upregulation (23.1-fold in the forebrain and 36.7-fold in the midbrain). *Fgf3* was also strongly upregulated in both brain regions (approximately five-fold), while *Fgf15* was only slightly upregulated in the midbrain (**Fig. 4C**). Interestingly, despite massive induction of *Fgf3* and *Fgf4* levels in the anterior forebrain, very few other genes were dysregulated at this stage (**Fig. S4C**). In contrast, the midbrain displayed more widespread dysregulation, which may explain the early neural expansion of the midbrain. In summary, our data revealed that deletion of the C1-C4 boundary fused the *Ano1* and FGF domains and led to strong ectopic expression of the FGF genes in the brain, which may result in expanded neural tissue and encephalocele. Supporting our hypothesis that increased FGF activity affects skull development, a previous study has revealed that induction of *Fgf3* and *Fgf4* driven by random insertion of viral elements can cause craniofacial dysmorphology (Carlton et al., 1998).

### Loss of CTCF-mediated insulation exposes FGF genes to distal brain enhancers of *Ano1*

The strong upregulation of the FGF genes and recapitulation of the brain *Ano1* expression pattern suggest that loss of the C1-C4 boundary causes ectopic contacts with enhancers located in the domain that harbors the *Ano1* gene. To identify candidate regulatory elements, we analyzed histone modification patterns in mouse wild-type midbrains at E11.5. The intronic region of *Ano1* showed two distinct H3K27ac peaks suggestive of putative active enhancers that may drive its expression in the brain (**Fig. 5A**, bottom panel). In line with its expression, *Fgf15* also had H3K27ac enrichment in the midbrain. Furthermore, we saw strong enrichment of the polycomb-deposited repressive mark H3K27me3 at *Fgf3* and *Fgf4* in wild-type brain, which likely contributes to their transcriptional silencing. We hypothesized that loss of insulation in embryos without the C1-C4 cluster could induce ectopic contacts of the FGF genes with these distal *Ano1* brain enhancers. To test this, we performed RCMC in micro-dissected midbrains at E11.5 to directly map the interaction of the distal enhancers with FGF genes, which could not be visualized at the resolution obtained from CHi-C experiments. As we saw with CHi-C data, we observed loss of both upstream and downstream CTCF-anchored loops (white arrowhead in **Fig. 5A**) due to deletion of the *C1-C4*^*flx*^ rescue cassette, and an increase in contacts between CTCF motifs of the clusters surrounding C1-C4. Surprisingly, control embryos with intact *C1-C4*^*flx/flx*^ rescue cassette had a weak but clearly visible interaction between *Fgf3* and the most centromeric of the *Ano1* intronic enhancers. However, the frequency of this interaction increased in *C1-C4*^*flx/flx*^*;Cre* mutant midbrains, where the C1-C4 motifs were deleted (black arrowhead in **Fig. 5A**). *Fgf15*, which has a very similar brain expression pattern to *Ano1*, also interacted with the *Ano1* brain enhancers. To confirm these interactions more precisely, we performed promoter capture Micro-C using the three FGF promoters as viewpoints (**Fig. 5B; S5A**). This strategy confirmed that deletion of the C1-C4 cluster resulted in loss of interactions at the CTCF boundary generated by the *C1-C4*^*fl*x^ rescue cassette (gray highlighted region on the right). There was also a sharp increase in long-range interactions of the *Fgf3* and *Fgf15* promoters specifically with the most centromeric of the *Ano1* intronic brain enhancers (highlighted region on the left). Rather than increased interactions specifically at this enhancer, *Fgf4* showed higher contact frequency across the entire *Ano1* domain (see subtraction track in **Fig. 5B** where the signal from *C1-C4*^*flx/flx*^ cells was subtracted from *C1-C4*^*flx/ flx*^*;CRE*). In summary, loss of CTCF-mediated insulation at the C1-C4 boundary promotes physical interaction of FGF promoters with putative *Ano1* brain enhancers.

**Figure 5.**
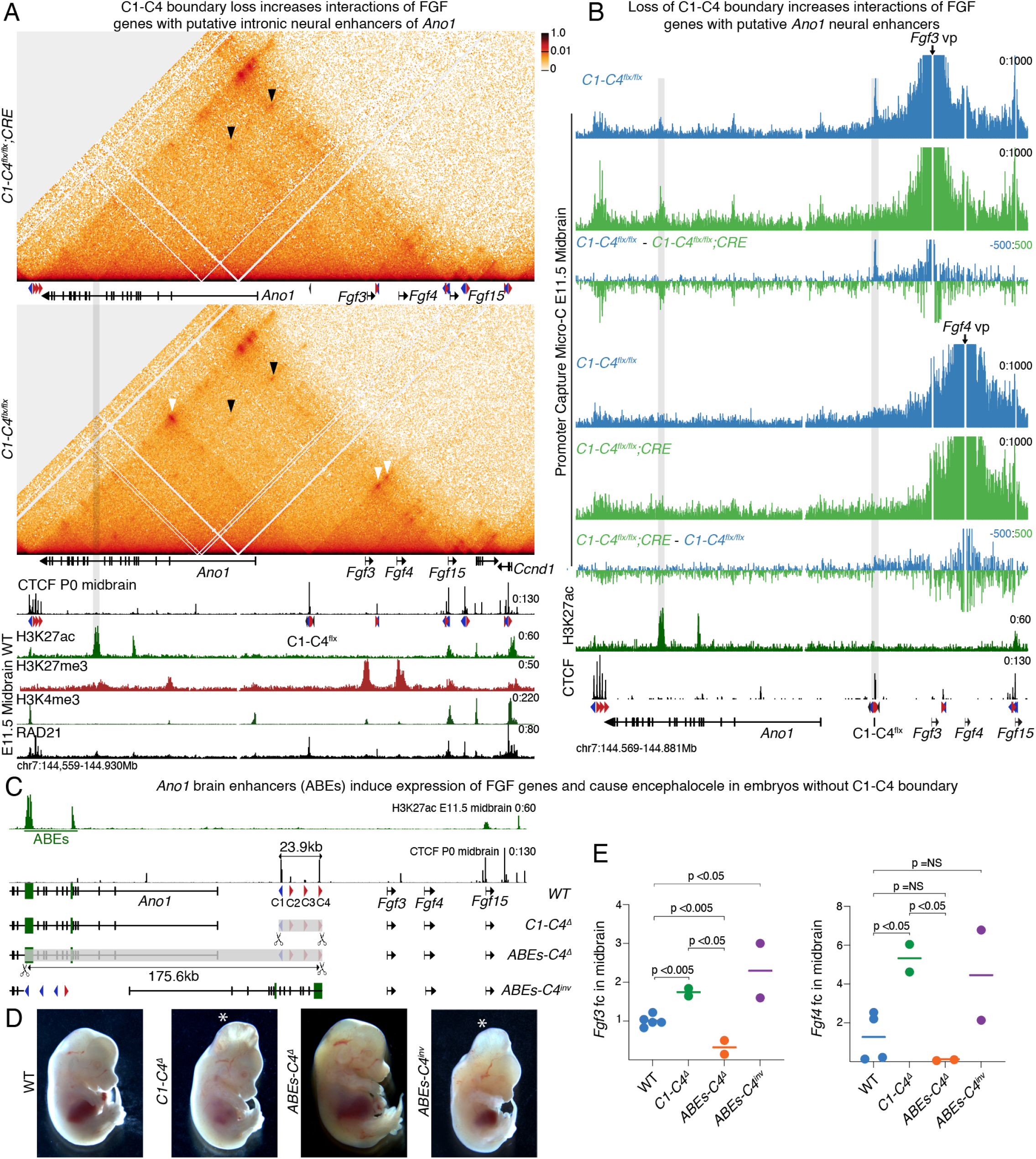
Loss of CTCF-mediated insulation exposes FGF genes to distal brain enhancers of *Ano1*. **A** Region capture micro-c (RCMC) 1D interaction frequency heatmap at 400bp resolution of *C1-C4*^*flx/flx*^;CRE and *C1-C4*^*flx/flx*^ microdissected E11.5 midbrains. White arrowhead shows loops between CTCF motifs of the C1-C4”^flx^ rescue cassette and the motifs in the surrounding clusters. Black arrowheads point to interactions between the most centromeric putative *Ano1* brain enhancer and *Fgf3* and *Fgf15*. All data, CHiP-seq shown below were mapped to a custom mm10 genome where the the C1-C4 cluster was replaced by the C1-C4^flx^ rescue cassette. Gray highlight shows the putative *Ano1* brain enhancer **B** Promoter Capture Micro-C shown at 50bp resolution for the *Fgf3* and *Fgf4* promoters. Gray highlight shows interactions between the FGF promoters and the *C1-C4*^*flx*^ rescue cassette or with the putative *Ano1* brain enhancer. **C** Schemes of alleles generated by zygotic electroporation to analyze phenotypes and gene expression at E13.5. **D** Representative images of embryos with each of the shown alleles (n= 6/6, 3/3, 2/2, 2/2). Asterisk indicates embryos with encephalocele. E qPCR measurement of *Fgf3* and *Fgf4* in E13.5 dissected midbrains using the ΔΔCT method and Gapdh as a reference. Each circle represents an embryo and the bar the median value for each genotype. A Wilcoxon two-sided test was performed to assess significance

We then did another genetic rescue experiment to prove that induction of the FGF genes—triggered by ectopic contacts with the distal putative *Ano1* brain enhancers (ABEs)— results in encephalocele. Specifically, we hypothesized that extending the 23.9kb deletion of the C1-C4 cluster to include these candidate intronic regulatory elements, would prevent the ectopic activation of the FGF genes—despite boundary deletion—and rescue the encephalocele phenotype. To test this, we electroporated zygotes with Cas9 and gRNAs designed to generate such deletion and analyzed phenotypes and gene expression directly in E13.5 founder embryos (**Fig. 5C**). As a positive control, we also re-targeted the CTCF cluster C1-C4 which, as expected, resulted in encephalocele in embryos carrying the *C1-C4*^*Δ*^ allele (**Fig. 5D**). Strikingly, all embryos with the larger 175.6kb deletion allele—that we named *ABEs-C4*^*Δ*^—displayed no overt phenotype at E13.5. We also screened for inversions where the targeted region in *ABEs-C4*^*Δ*^ was inverted and repositioned the ABEs closer to FGF genes, putting the CTCF boundary further away towards the centromeric side. In contrast to embryos where the enhancers were deleted, the encephalocele phenotype was clearly visible in all *ABEs-C4*^*inv*^ embryos (**Fig. 5D**). In line with the rescue of the encephalocele phenotype, *Fgf3* and *Fgf4* levels were significantly reduced in the midbrain of *ABEs-C4*^*Δ*^ mutants as compared to wildtype littermates or *C1-C4*^*Δ*^ mutants (**Fig. 5E**). *ABEs-C4*^*inv*^ mutants showed higher levels of *Fgf3*, likely because the inversion allele places the ABEs closer to the FGF genes, as observed for other loci (Zuin et al., 2022). Together, these data reveal that the loss of the C1-C4 boundary exposes the three FGF genes to brain enhancers of *Ano1*, which results in strong ectopic brain expression and encephalocele.

### Deletion of a single CTCF motif within a large multi-motif cluster can compromise its insulator function

Our data show that the chromatin domain boundary established by the C1-C4 cluster has a remarkable ability to completely insulate the *Ano1* brain enhancers from activating the FGF genes in wild-type embryos. This is especially clear for *Fgf4* whose expression is almost undetectable before recombination but becomes highly upregulated once the C1-C4 cluster is deleted. To elucidate the contribution of the different types of CTCF motifs within the C1-C4 cluster to insulate the FGF genes, we targeted the boundary by injecting zygotes with Cas9 and gRNAs in different combinations (**Fig. 6A**). The C1 motif is in reverse orientation, pointing to the domain of the *Ano1* gene, whereas the C2-C4 motifs are in the forward orientation, towards the domain with the three FGF genes. To understand if all four CTCF motifs are required to segregate the FGF genes from the ABEs, and if their orientation determines different insulation strengths, we generated two deletion mutants—*C1*^*Δ*^ and *C2-C4*^*Δ*^ (**Fig. 6B**). Surprisingly, all E13.5 embryos with just the *C1*^*Δ*^ deletion recapitulated the encephalocele phenotype observed in *C1-C4*^*Δ*^, while *C2-C4*^*Δ*^ mutants did not display any phenotype. Notably, the expression levels of *Fgf3* and *Fgf4* in *C1*^*Δ*^ mutants were significantly upregulated and comparable to the full cluster deletion of *C1-C4*^*Δ*^ mutants (**Fig. 6C**). In contrast, *C2-C4*^*Δ*^ mutant embryos had no statistically significant differences in *Fgf3 and Fgf4* midbrain expression compared to wild-type littermates. In summary, these data highlight the importance of insulation by the C1-C4 cluster to ensure that *Fgf3* and *Fgf4* are not expressed in the midbrain and how loss of a single motif within a multi-motif cluster can compromise its insulator function and result in ectopic expression of these genes. Our data also suggest that CTCF motifs oriented towards the side of active enhancers may be better transcriptional insulators than those oriented to inactive chromatin regions

**Figure 6.**
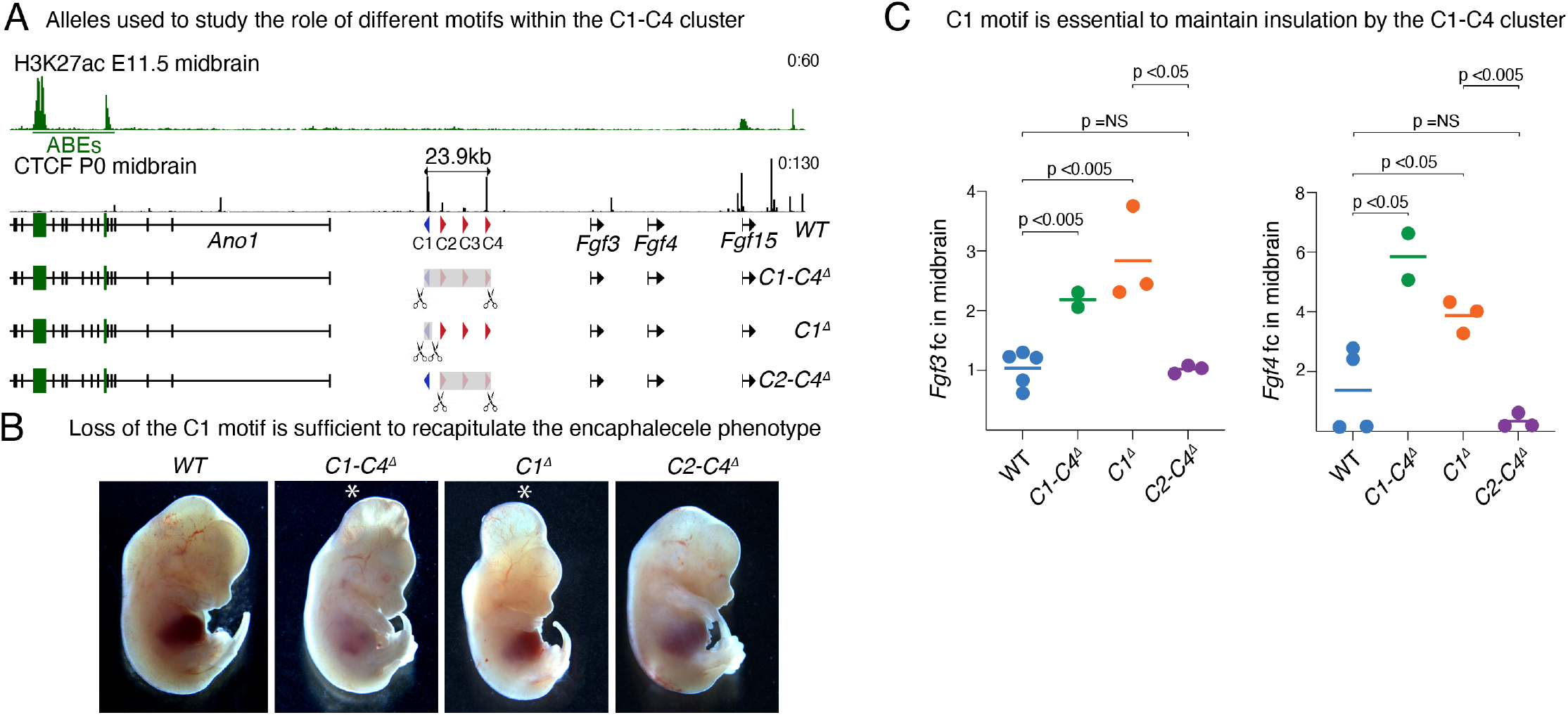
Loss of a single motif within C1-C4 recapitulates deletion of entire C1-C4 cluster. **A** Schemes of alleles generated by zygotic electroporation to analyze phenotypes and gene expression at E13.5. **B** Representative images of embryos with each of the shown alleles (n= 6/6,3/3, 6/7,3/3). C qPCR measurement of *Fgf3* and *Fgf4* in E13.5 dissected midbrains using the Δ ΔCT method and *Gapdh* as a reference. Each circle represents one embryo and the bar the median value for each genotype. A Wilcoxon two-sided test was performed to assess statistical significance.

## DISCUSSION

The discovery that vertebrate genomes fold into chromatin domains delimited by CTCF binding presents an attractive and simple model to understand how the spatial range of enhancers can be restricted to genes within the same domain. However, disruption of these structures often results in minor changes in gene expression, and modest phenotypes (Paliou et al., 2019; Rodriguez-Carballo et al., 2017; Sarro et al., 2018). This raises doubts on the physiological impact of chromatin domains and their roles in gene regulation. Here, we show that in some developmental contexts, such impact can be dramatic. In the developing murine brain, although *Fgf3* and *Fgf4* are polycomb targets marked for silencing by H3K27me3 enrichment, loss of their centromeric domain boundary induces strong ectopic upregulation by the intronic enhancers of *Ano1*, leading to orofacial clefts, encephalocele, and fully penetrant perinatal lethality. Strikingly, loss of just one CTCF motif—pointing towards the *Ano1* enhancers—is sufficient to recapitulate these severe phenotypes.

We hypothesized that disruption of domains with multiple developmental regulators may induce particularly striking changes in gene expression and animal development. While this was true when we targeted the centromeric domain boundary and observed prominent defects in brain development, this phenomenon was not manifested in other developmental processes controlled by the FGF genes nor when we targeted the CTCF clusters separating them. For example, we showed that although blastocyst development requires precise control of FGF4 levels, CTCF boundaries around this gene are not necessary for early development. Our data suggest that these different phenotypic outcomes are caused by different enhancer-specificities of the FGF promoters. In the brain, all three FGF genes have strong compatibility with a distal enhancer of *Ano1* and therefore CTCF insulation is essential to prevent their ectopic activation. In contrast, in the blastocyst these same genes are remarkably specific to their own proximal enhancers and gene regulation is independent of CTCF-mediated boundaries. As explained below, we propose that different strategies to achieve specificity in enhancer-promoter interactions and gene expression may be driven by how insulation and domains are built during development.

Mammalian fertilization is characterized by sequential reestablishment of chromatin structure. Domain boundaries are initially very permeable and gain insulation strength with each cleavage stage (Borsos et al., 2019; Du et al., 2017; Flyamer et al., 2017; Ke et al., 2017). ES cells, which mimic the epiblast of the blastocyst, also show weaker boundaries than more differentiated cells such as neurons or even neural progenitors (Bonev et al., 2017). Perhaps early in development, gene regulation must rely on more stringent promoter specificity that may be better conferred by proximal enhancers. In contrast, once domain boundaries provide better insulation, cell-type specific expression can be driven by more distal enhancers, whose spatial range of gene activation is controlled by CTCF-mediated boundaries. In line with this, depletion of maternal and zygotic CTCF does not cause dramatic dysregulation of gene expression in early blastocysts, providing further support to the hypotheses that early in development, CTCF is less important to ensure enhancer–promoter specificity (Andreu et al., 2022).

The chromatin domain that contains *Ano1*, the three FGF genes, and *Ccnd1* is in a syntenic region found across vertebrates. Importantly, clusters of CTCF motifs between these genes are also conserved in most vertebrate species. Interestingly, duplication of this region in the genome of Ridgeback dogs leads to dermoid sinus and causes dysregulation of FGF genes (Salmon Hillbertz et al., 2007). In humans, duplication of the chromosomal 11q13 region, which includes *FGF3* and *FGF4* causes intellectual disability, craniosynostosis and microcephaly (Grillo et al., 2014; Jehee et al., 2007; Martinez Anaya et al., 2023; Ziebart et al., 2013). Furthermore, global DNA hypermethylation seen in gastrointestinal stromal tumor (GIST) patients leads to recurrent loss of CTCF binding in the boundary between *ANO1* and the FGF genes, and higher levels of FGF activity (Flavahan et al., 2019). Although it isn’t yet clear how much the structure of the chromatin domain is affected by these different chromosomal rearrangements, these observations suggest that spatial organization and regulatory interactions within this region may be preserved throughout evolution to fulfill essential developmental functions.

In contrast to mouse development where the loss of a single CTCF motif—if oriented towards an active enhancer—was sufficient to activate ectopic FGF expression, in human GIST cell lines, disruption of all four CTCF motifs found in its cluster is required for induction of *FGF3* by *ANO1* (Kim et al., 2024). The difference in sensitivity between the two models may be related to the boundary, or to the enhancers that drive expression of *Ano1* in each of these two tissues, i.e. the mouse midbrain, and the human intestine. Nonetheless, our work, together with these observations across multiple vertebrates, suggests that structural perturbation of highly conserved chromosomal domains may be more likely to result in strong gene dysregulation and more severely affect physiological processes (Fudenberg and Pollard, 2019; Vietri Rudan et al., 2015). Yet, this will not be true in all contexts. Many factors, such as enhancer type and strength, orientation and placement of CTCF motifs within a cluster, frequency and mediators of enhancer–promoter interactions, among others, are likely to affect physiological outcomes and much work is still needed to understand their impact.

Recent sequencing efforts have highlighted the high prevalence of sequence variants in the non-coding genome. Our work further emphasizes the importance of whole genome sequencing to consider how even small changes in CTCF binding motifs can mis-wire enhancer–promoter interactions, induce ectopic gene expression, and initiate disease states or perturb development. Our data also provide important insight into potential congenital origins of encephalocele, and it will be important to investigate whether human encephalocele patients present sequence variants that may affect CTCF binding at the boundary between *ANO1* and the FGF genes.

## METHODS

### Animals and transgenic line generation

Zygotic injection of Cas9-gRNA ribonucleoproteins was used for mouse line generation, as previously described (Chakraborty et al., 2023). sgRNAs were designed using sgRNA Scorer 2.0 (Chari et al., 2017) and subsequently tested for editing activity in P19 cells, as previously described (Gooden et al., 2021). The best gRNAs were then purchased as synthetically modified RNAs (Synthego). Cas9 protein was generated in house (Protein Expression Lab, Frederick National Lab) using an *E. coli* expression plasmid obtained as a gift from Niels Geijsen (Addgene #62731) (D’Astolfo et al., 2015). Superovulated C57Bl6/6NCr female mice were used as embryo donors. Zygotes were collected on day E0.5, microinjected and allowed to recover for 2 hours by incubation (5% CO2, 37°C), following which, viable embryos were surgically transferred to oviducts of pseudopregnant recipient females. The cocktail used for injection comprised of 75 ng/μl of in vitro synthetized gRNAs (Synthego) and 50ng/ μl of Cas9 protein in 50μl total volume, kept on dry ice until just prior to microinjection. For insertion of a repair template, 75 ng/μl of single-stranded DNA was added to microinjection cocktail. Founder mice were then bred to C57Bl6 mice from The Jackson Laboratory. Repair templates, gRNAs, primers used for genotyping, and sequencing of ligation junctions of the deletion lines can be found in **Table S1**. Targeting for analysis of founder F0 E14.5 embryos was done by zygotic electroporation using the Bex zygote genome editing electroporator and the following conditions: 30V, 1ms Pd on, 1000ms pd off. For these experiments, 12μl of 250ng/ μl of Cas9 (IDT 1081058), 100ng/μl of gRNA (ordered from IDT). The following Cre lines were obtained from the Jackson Laboratory: E2A-CRE (strain 003724) (Lakso et al., 1996), ROSA26-Cre^ERT2^ (strain 008463) (Ventura et al., 2007), Wnt1-Cre (strain 022501) (Lewis et al., 2013), and Sox10-Cre (strain 025807) (Matsuoka et al., 2005). Mouse studies were performed according to NIH and PHS guidelines and only after protocols were approved by the Animal Care and Use Committees of the National Cancer Institute and Eunice Kennedy Shriver National Institute of Child Health and Human Development. NCI-Frederick, where generation of transgenic mouse lines was made, is accredited by AAALAC International and follows the Public Health Service Policy for the Care and Use of Laboratory Animals. Animal care was provided in accordance with the procedures outlined in the “Guide for Care and Use of Laboratory Animals (National Research Council; 1996; National Academy Press; Washington, D.C.).

### Skeletal stainings

Skeletal analyses were performed using Alcian blue and Alizarin red staining method as described previously (Mallo and Brandlin, 1997). Briefly, fetuses were obtained at E18.5 by caesarean section, eviscerated and soaked in water for 2-4 hours. After a one-minute heat shock at 65°C fetuses were skinned and fixed in 100% ethanol for 2 days. Fixed fetuses were incubated with 150mg/l Alcian blue in 80% ethanol and 20% acetic acid for 12 hours followed by overnight incubation in 100% ethanol. Fetuses were treated with 2% KOH for 6 hours and stained with 50mg/l Alizarin red in 2% KOH for 3 hours and cleared again in 2% KOH for 12-20 hours. Stained and cleared fetuses were stored in 25% glycerol in water before imaging.

### Micro-CT scans

This was done by the Mouse Biology Program, University of California, Davis, CA, United States as previously described (Shankar et al., 2022). E18.5 embryos were incubated in a hydrogel stabilizing solution (4% PFA, 4% acrylamide, 0.05% bis-acrylamide, 0.25% VA044 Initiator, 0.05% saponin in PBS) for three days at 4°C to preserve tissue integrity. Thereafter, the vials containing the embryos were placed in a desiccation chamber and saturated with nitrogen gas to replace the air and embryos were incubated in a 37°C water bath. Finally, embryos were removed from the encasing hydrogel, swiped clean and immersed in a Lugol solution [0.7% iodine solution (0.1N)] for at least 24 hours at room temperature while rocking and then oriented and embedded in 1% agarose and oriented for μCT imaging. Mouse embryos imaged at the Center for Molecular and Genomic Imaging (UC Davis) with high-resolution X-ray CT. Three embryos were embedded stacked in agar in a in a standard test tube that fit the embryos tightly and the conical vial was glued to a sample base for imaging. Embryos were imaged using a high resolution MicroXCT-200 specimen CT scanner (Carl Zeiss X-ray Microscopy). The CT scanner has a variable x-ray source capable of a voltage range of 20-90kV with 1-8W of power. Embryos were placed on the scanner’s sample stage, which has a submicron level of position adjustments. Scan parameters were adjusted based on the manufacturers recommended guidelines. The systems 0.4x detector was used for imaging. The source and detector distances were adjusted based so that the Detector-RA and Source-RA distances were 122.5mm and 25mm respectively. Once the source and detector settings were established, the optimal x-ray filtration was determined by selecting among one of 12 proprietary filters: LE3 was the filtration selected. Following this procedure, the optimal voltage and power settings were determined for optimal contrast (40kV and 200microAmp). 1,600 image projections were obtained over a 360-degree rotation. The camera pixels were binned by two to increase signal to noise in the image and the source-detector configuration resulted in a voxel size of 11.4791 microns. Images were reconstructed with a smoothing factor of 0.7 and a beam hardening of 0.2 into 16-bit values with common global minimum and maximum values for proper histogram matching.

### ES establishment and differentiation

Mouse embryonic stem cells lines were derived and genotyped as previously described in (Chakraborty et al., 2023). For assays with cells in the Epiblast-like state, cells were grown in serum-free 2i media constituted of Neurobasal medium (Thermo Fisher, 21103049), DMEM/ F12 Nutrient mixture (Thermo Fisher, 21103049), 1% penicillin—streptomycin (Thermo Fisher, #15140163), 2 mM Glutamax (Thermo Fisher, #35050079), β-mercaptoethanol supplemented with N2 (Thermo Fisher, #17502001), B-27 (Thermo Fisher, #17504001), MEK/ERK pathway inhibitor (PD0325901, Reprocell, #04-0006-02), GSK3 signaling inhibitor (CHIR99021, Reprocell, #04-0004-02) and leukemia inhibition factor (LIF, Sigma, #ESG1107). Before differentiation, ES cells were grown on mouse embryonic fibroblasts and serum containing media: Knockout Dulbecco’s Modified Eagle Medium (DMEM) (Thermo Fisher, #10829018) with 15% FBS (VWR, #97068-091), 2 mM Glutamax (Thermo Fisher, #35050079), 0.1 mM β-mercaptoethanol, 0.1 mM MEM non-essential amino acids (Thermo Fisher, #11140050), 1 mM sodium pyruvate (Thermo Fisher, #11360070), 1% penicillin—streptomycin (Thermo Fisher, #15140163) and Recombinant mouse LIF (Sigma, #ESG1107). Differentiation into the primitive endoderm-like state (XEN cells) was done as previously described (Niakan et al., 2013). Briefly, ES cell lines were first grown on mouse embryonic fibroblasts, enzymatically passaged with 0.05% trypsin and feeder depleted for 45 mins. Cells were washed twice with standard XEN media: advanced RPMI (Thermo Fisher, #12633012) with 15% FBS (VWR, #97068-091), 1% penicillin—streptomycin (Thermo Fisher, #15140163) and 0.1 mM β-mercaptoethanol to remove residual LIF. Cells were counted, 96000 cells were plated in gelatinized 6-well plates and grown overnight in standard XEN media. After 24 hours, media was replaced with standard XEN media supplemented with 0.01μM retinoic acid dissolved in DMSO, 10ng/ml activin, 24ng/ml recombinant FGF2 and 1μg/ml heparin. Cells were grown in presence of supplements for 2-3 days depending on morphology and confluency and media was replenished every 24 hours. Cells were passaged and grown for 7-10 days until XEN like colonies with stellate and refractile morphologies emerged. XEN-like colonies were scraped and picked with a pipette under the microscope and plated in gelatinized plates to enrich their populations and cultured for an additional 2 weeks until cell morphology was homogenous.

### RNA-seq

RNA from the developing brain was isolated using trizol reagent. After confirming that the RNA integrity number for each sample was above 8, libraries were prepared using TruSeq Stranded mRNA prep kit with PolyA purification and sequenced on Novaseq6000. For RNA-seq in single blastocysts, DNA and RNA was isolated from each embryo using Dynabeads mRNA DIRECT Purification Kit (Thermo Fisher, 61012) as previously described (Huffman et al., 2012). Blastocysts at E4.5 were flushed out of the uterine horns from super-ovulated females using M2 media and transferred into 50 μl of pre-warmed lysis buffer (100mM Tris-HCl pH-7.5, 500mM LiCl, 10mM EDTA pH-8, 1%LiDS, 5mM DTT) in DNA low binding PCR tubes. Lysed embryos were stored in -20C and processed within a few weeks. Dynabeads Oligo(dT)_25_ mRNA isolation beads were warmed to room temperature for 30 mins and rinsed with 100 μl of lysis buffer by vortexing continuously for 5 mins. 10 μl of bead suspension was used per embryo lysate. The poly-A tail of mRNA was allowed to anneal to beads by mixing the embryo lysates and bead suspension in a vortexer for 5 mins at low speed, followed by 5 mins incubation without shaking. Tubes were briefly spun and placed in a magnetic stand to collect the clear supernatant, which was used for DNA isolation using 2X SPRI beads to determine the genotype of each embryo. mRNA bead complexes were washed twice using Wash Buffer A and twice with Wash Buffer B by vortexing for 5 mins each. cDNA and library prep were then processed using an adaptation of the smartseq2 protocol (Picelli et al., 2014; Satija et al., 2015; Trombetta et al., 2014). Buffers were removed and beads annealed with mRNA was resuspended in 12.5μl of RNA suspension mix containing 20U/μl SUPERase-In RNase inhibitor (Thermo Fisher, #AM2694), 10mM each dNTP Mix (New England Biolabs, #N0447L) and 100μM polydT oligonucleotide primer (custom sequence AGACGTGTGCTCTTCCGATCTTTTTTTTTT TTTTTTTTTTTTTTTTTTTTTVN synthesized by Integrated DNA Technologies). Bead suspension is heated at 75°C for 5mins to denature the RNA and supernatant is transferred into a new RNase free 0.2ml PCR tube. To each sample tube 7.5μl of first strand reverse transcription mix with 100mM DTT, 5X SuperScript IV Buffer, SuperScript™ IV Reverse Transcriptase (Thermo Fisher, #18090050), 20U/μl SUPERase-In RNase inhibitor (Thermo Fisher, #AM2694), 100μM of template switching oligo with 3G’s and an adaptor sequence same as polydT oligonucleotide (custom sequence ‘AGACGTGTGCTCTTCCGATCTNNNNNrGrGrG’, Integrated DNA Technologies) was added and incubated at 50°C for 60 mins to prepare the first strand of cDNA, followed by 85°C for 5mins to inactivate the enzyme. The first strand of cDNA is bound by the template switching oligo and a complementary sequence to the template switching oligo was synthesized, where a PCR oligo is hybridized (custom sequence ‘AGACGTGTGCTCTTCCGATCT’, Integrated DNA Technologies) and amplified. PCR cDNA amplification was done for 16 cycles (98°C for 15 secs, 67°C for 20 secs and 72°C for 10 mins) in a thermal cycler with KAPA Hifi HotStart ReadyMix (Roche, #9420398001). Pre-amplication cDNA mix was purified with 0.8X SPRI beads and quality of cDNA was assessed with a D5000 Screen Tape assay (Agilent, #5067-5588) using Agilent 4150 Tapestation. cDNA mix was diluted to 0.2ng/μl and used for library preparation as described earlier (Trombetta et al., 2014) using Nextera XT DNA Library Prep Kit (Illumina, #FC-131-1024). Briefly, in a 384 well plate, 1 μl of cDNA, 2μl of Tagment DNA Buffer and 1μl of Amplicon Tagment Mix was mixed in ice and tagmentation was done in a thermocycler at 55°C for 10 mins. Reaction was neutralized with 1 μl of Neutralize Tagment Buffer, incubated at room temperature for 5 mins and transferred to ice. Amplification PCR was set up with 3μl of Nextera PCR Mastermix and 2μl of each adaptor from Illumina DNA/RNA UD Indexes Set A (Illumina, #20027213) per sample. Tagmented DNA was incubated at 72°C for 3 mins, 95°C for 30 secs and 12 cycles of 95°C for 10 secs, 55°C for 30 secs, 72°C for 1 min and a final extension at 72°C for 5 mins. Library DNA was purified with 0.9X SPRI and quality was assessed using D1000 Screen Tape assay (Agilent, #5067-5585) before sequencing.

### RNA-seq analyses

RNA-seq analysis for brain was performed on 50-bp paired-end raw sequence reads using lcdb-wf v1.9rc [github. com/lcdb/lcdb-wf]. Adapters were trimmed, along with light quality trimming, using cutadapt v3.4 with parameters “-a AGATCGGAAGAGCACACGTCTGAACTCCAGTCA -q 20 --minimum-length = 25” (Martin, 2011). Sequencing quality was assessed with fastQC v0.11.9 (bioinformatics.babraham. ac.uk/projects/fastqc/) and evaluation for common sequencing contaminants was performed using fastqc_screen v0.14.0 with parameters “--subset 100000 --aligner bowtie2”. No significant quality issues were detected. Trimmed reads were mapped to the *Mus musculus* reference genome (GENCODE m18) using STAR v2.7.8a (Dobin et al., 2013) in one-pass mode with standard parameters from ENCODE long-RNA-seq pipeline. Aligned reads were counted using featureCounts from the subread package v2.0.1 (Liao et al., 2014) using default parameters, which excludes multimapping reads by default. Differential expression analysis on raw counts was performed with DESeq2 v1.22.1 (Love et al., 2014). A separate DESeqDataSet object was created for each brain region and the model included a blocking factor for sex using the model ∼sex + genotype. The genotype contrast was extracted followed by log2 fold change shrinkage using the ‘normal’ method. The analysis used DESeq2’s default of considering a gene differentially expressed if the false discovery rate (FDR) of the gene was <0.1. No log2 fold change threshold was applied. Count data were normalized by DESeq2-calculated sizeFactors for visualization purposes. RNA-seq analysis for blastocysts was similar to above except that in the upstream processing, the cutadapt parameters were changed to “-a CTGTCTCTTATACACATCT” and “-A CTGTCTCTTATACACATCT” to reflect the different library prep method. In DESeq2 a separate DESeqDataSet object was created for each deletion (along with matched control) using the model ∼genotype and the genotype contrast was extracted.

### Hybridization Chain Reaction (HCR)

HCR was performed as previously described (Anderson et al., 2020). Briefly, embryos were fixed in 4% PFA overnight at 4°C while rocking, then gradually dehydrated into 100% methanol where they can be stored indefinitely. Before beginning HCR, embryos are rehydrated and washed into PBT (PBS + 0.1% Tween), bleached with 6% hydrogen peroxide in PBS for 30 minutes, then digested with 10μg/mL ProK (Sigma 3115836001) in PBT for 12 minutes. Digestion was stopped by briefly washing with PBT then post-fixing in 4% PFA for 20 minutes at room temperature. Embryos are equilibrated in hybridization buffer at 37°C then hybridized with overnight at 37°C with probes for *Fgf3, Fgf4, Fgf15*, and *Ano1*, with the initiators B3, B4, B1, and B2 respectively. Hairpins were diluted in AMP buffer and rocked overnight at RT. Hairpins used for each initiator were: B3-546, B4-647, B1-488, B2-750. Embryos were then washed in PBT then PBTx (PBS + 0.1% TritonX-100) with DAPI for 36-48 hours at RT. Probes, hairpins, hybridization buffer, probe wash buffer, and amplification buffer were purchased from Molecular Instruments. Embryos were mounted in 1% ultra-low-melt agarose (Sigma A5030) then cleared in Ce3D++. All images of HCR-stained embryos were taken on a Nikon A1 confocal microscope using a Plan Apo 10x objective (NA: 0.45).

### Capture Hi-C and Promoter Capture Hi-C

Hi-C Libraries used for Capture and Promoter Capture were generated as previously described ^36717694^ with a few modifications. Briefly, for mouse ES cell lines, 1 million cells per sample were trypsinized, washed in growth media and fixed for 35 minutes at room temperature while rotating in 1mg/ml DSG (D2650) in 1 ml of PBS. 1% formaldehyde (Thermo 28908) was then added, and samples fixed for 10 more minutes. CHiC was processed separately for the two independent lines of each genotype that had been established from two independent blastocysts. E11.5 midbrains were dissected in chilled artificial cerebrospinal fluid (ACSF: 87 mM NaCl, 26 mM NaHCO3, 2.5 mM KCl, 1.25 mM NaH2PO4, 0.5 mM CaCl2, 7 mM MgCl2, 10 mM glucose, 75 mM sucrose, saturated with 95% O2, 5% CO2, pH 7.4). Cells were then incubated with 0.1% Pronase in PBS for 15 minutes (Sigma 10165921001) at room temperature. Cells were then fixed as described above. To stop fixations, Glycine was added at final concentration of 0.13M and incubated for 5 minutes at RT and 15 minutes on ice. Cells were then washed once in cold PBS, centrifuged at 2500g 4°C for 5 mins (these centrifugation conditions were used for all washes following fixation) and pellets frozen at -80°C. Thawed cell pellets were then lysed (10mM Tris-HCL pH8, 10mM NaCl, 0.2% Igepal CA-630, Roche Complete EDTA-free Sigma #11836170001) and digested with 600U of DpnII (NEB). Biotin fill-in was done by incubating cells with a mixture of dCTP, dTTP, dGTP, Klenow polymerase (NEB M0210L) and 37.5μl Biotin-14-dATP (Thermo 19524016) for 4h at RT while shaking at 900rpm for 10 seconds every 5 minutes. Ligation was done overnight at 16°C using T4 ligase (NEB cat #M0202M). Sonication was done using Covaris onetube-10 AFA strips using the following parameters for a 300bp fragment size (Duration: 10secs, repeat for 12 times, total time 120 secs, peak power-20W, duty factor 40%, CPB-50). Library-prep with material on the beads was done using the Hyper Prep kit (Roche KK8502).

For Capture Hi-C capture reactions, 1ug of Hi-C library per sample were hybridized with 120 nucleotide biotinylated RNA probes using the SureSelect kit from Agilent and PCR amplification using the polymerase from the Kapa Hyper Prep Kit. For promoter Capture Hi-C, 1μg of Hi-C library per sample was hybridized with 120bp biotinylated oligos using the SeqCap EZ kits (Roche). Following washes, material was amplified by PCR using the Kapa polymerase from the Hyper Prep kit (Roche KK8502). Material from different samples was then combined and 1μg of pooled libraries was recaptured. Probes used for these assays can be found in **Table S1**. omains delimited by CTCF binding presents an attractive and simple model to understand how the spatial range of enhancers can be restricted to genes within the same domain. However, disruption of these structures often results in minor changes in gene expression, and modest phenotypes (Paliou et al., 2019; Rodriguez-Carballo et al., 2017; Sarro et al., 2018). This raises doubts on the physiological impact of chromatin domains and their roles in gene regulation. Here, we show that in some developmental contexts, such impact can be dramatic. In the developing murine brain, although *Fgf3* and *Fgf4* are polycomb targets marked for silencing by H3K27me3 enrichment, loss of their centromeric domain boundary induces strong ectopic upregulation by the intronic enhancers of *Ano1*, leading to orofacial clefts, encephalocele, and fully penetrant perinatal lethality. Strikingly, loss of just one CTCF motif—pointing towards the *Ano1* enhancers—is sufficient to recapitulate these severe phenotypes.

### Region Capture Micro-C

Libraries for Micro-C were prepared as previously described (Brown et al., 2023; Slobodyanyuk et al., 2022) with a few changes as detailed below. Dissection, dissociation, and fixation of cells grown in vitro and from E11.5 midbrains was done as described for Hi-C libraries. MNase was titrated with 1 million cells at 3U, 5U and 10U for 20 min at 37 °C in 100μl. Then, the micrococcal nuclease reaction was carried out with 5 million cells per replicate, and the micrococcal nuclease step was scaled up to 500 μL with the optimal concentration of micrococcal nuclease determined with titration experiment. Phosphorylation of DNA ends was done for 30 minutes at 37 °C using T4 Polynucleotide Kinase (NEB M0201), 3’ exonuclease activity of Klenow polymerase (NEB M0210) was done also at 37 °C for 30 minutes while shaking. Biotin incorporation with dATP and dCTP (Jena Biosciences NU-835-BIO14-L and NU-809-BIOX-L) was done at 25 °C for 90 minutes. T4-mediated proximity ligation was always done overnight with T4 ligase (NEB M0202M). Proteinase K (NEB P8107S) incubation was also performed overnight at 65 °C. After phenol/chloroform/iso-amyl alcohol extraction, the sample was split into two equal aliquots and was purified on two Zymo DNA clean and concentrator kit columns, eluted with 25 μL (preheated to 70 °C) elution buffer then pooled. Samples were loaded onto 3% TBE NuSieve GTG agarose gel in four separate wells. After excision of fragments containing dinucleosomes from each lane (>220bp, <400bp), samples were purified using a Zymo Gel DNA Recovery kit to extract DNA. DNA fragments were not polished prior to streptavidin binding. For biotin purification, 50 μL of Streptavidin C1 beads (Thermo 65002) were used per sample. Libraries were prepared using the KAPA Biosystems HyperPrep kit (Roche KK8502) and Illumina primers. After running a small-scale PCR to determine the optimal number of cycles for the required yield, four 50μL PCR reactions were set up for each sample as instructed by manufacturers. 200 μL was transferred to a new tube and incubated with 0.9x SPRI beads. Samples were sequenced by 50 bp pair-end sequencing with a NovaSeq 6000 with an SP100 kit by the NICHD Molecular Genomics Core. Region capture of Micro-C samples was done as described previously (Goel et al., 2023) and as recommended by the Twist Bioscience’s Standard Hybridization Target Enrichment Protocol with a few modifications. Specifically, to increase stringency, all wash steps were done at 70 °C instead of 48 °C and room temperature. The regions covered by probes can be found in **TableS1**. While biotinylated probes used in cells were of 120bp, the probes used in embryonic material were of 80bp.

### Region Capture Micro-C and CHIC analysis

Capture Hi-C and Region Capture Micro-C data were mapped to the mm10 genome using BWA (0.7.17) (Li and Durbin, 2009). BAM files were then processed to pairs format, filtered, sorted, selected for pairs that included both ends of an interaction within the captured region, and deduplicated using pairtools (1.0.2) (Open2C et al., 2023). Technical replicates resulting from sequencing of the same sample more than once were merged prior to duplicate removal. Replicates where libraries were obtained from different cells were merged post duplicate removal. Cool and balanced mcool files were generated using cooler (0.8.11) (Abdennur and Mirny, 2020). Visualization of contact data as heatmaps was done using higlass (Kerpedjiev et al., 2018) at resgen. io. For generation of viewpoint tracks, pairtools select was used to identify pairs where one mate is located within the viewpoint of interest. A custom genome containing the floxed C1-C4 rescue cassette instead of the C1-C4 boundary was generated using reform (github.com/gencorefacility/reform) to create a modified fasta mm10 genome file and GTF file. These files were then used instead of mm10.fa for alignment

### Cut&Run

Cut&Run was done as previously described (Thompson et al., 2022). Briefly, ES cells were detached from culture plates using Accutase (Sigma), counted and 200,000 cells per clone were spun at 600g for 3 mins at room temperature. Supernatant was discarded and cells were resuspended in Wash Buffer with 20mM HEPES pH 7.5, 150mM NaCl, 0.5mM Spermidine and 1x Protease inhibitor cocktail at 600g for 3 mins at room temperature. BioMag Plus Concanavalin A beads (Bangs Laboratories) were equilibrated in Binding Buffer with 20 mM HEPES pH 7.5, 10 mM KCl, 1 mM CaCl_2_ and1 mM MnCl_2_. Cell pellets were resuspended in Wash Buffer, mixed with a slurry of equilibrated Concavalin A coated magnetic beads, and rotated for 10 mins at room temperature. For each sample, 10 μl bead slurry was used and were placed on a magnetic separator to discard the supernatant. Beads were again resuspended in Wash Buffer containing 2 mM EDTA, 0.1% bovine serum albumin, 0.05% Digitonin, and 1:50 dilution of primary antibody against H3K27Ac (Abcam, ab4729). This was incubated on a nutating platform for 2 hours at room temperature. After incubation, beads were washed twice in Digitonin Buffer (20 mM HEPES pH 7.5, 150 mM NaCl, 0.5 mM Spermidine, 1x Roche Complete Protease Inhibitor no EDTA, 0.05% Digitonin and 0.1% bovine serum albumin), then incubated with lab prepared pA-MNase (600 μg/ml, 1:200) in Digitonin Buffer for 1 hour at 4 °C. After incubation, beads were washed twice, resuspended in 150 μl of Digitonin Buffer, and equilibrated to 0°C before adding 2mM CaCl_2_ and incubated at 0°C for 1 hour. After incubation, 150 μl of 2X Stop Buffer containing 200 mM NaCl, 20 mM EDTA, 4 mM EGTA, 50 μg/ml RNase A and 40 μg/ml glycogen was added. Beads were incubated for 30 mins at 37°C and then spun at 16,000 g for 5 mins at 4°C. Supernatant was transferred, mixed with 3 μl 10% SDS and 1.8U Proteinase K (NEB #P8107S) and incubated overnight at 55°C, shaking at 900 rpm. After incubation, 300 μl of 25:24:1 Phenol/Chloroform/ Isoamyl Alcohol was added, solutions were vortexed, and transferred to Maxtrack phase-lock tubes (Qiagen #129046). Tubes were centrifuged at 16,000 g for 3 mins at room temperature. 300 μl of Chloroform was added, solutions were mixed by inversion, and centrifuged at 16,000 g for 3 mins at room temperature. Aqueous layers were transferred to new tubes and DNA was isolated by ethanol precipitation and resuspended in 10 mM Tris-HCl pH 8.0 (Thermo Fisher #15568025). Cut&Run libraries were prepared following the SMARTer ThruPlex TAKARA Library Prep kit with small modifications. For each sample, 10 μl of double stranded DNA, 2 μl of Template Preparation D Buffer and 1 μl of Template Preparation D Enzyme were combined, and End Repair and A-tailing was performed in a thermocycler with a heated lid at 22°C, for 25 mins and 55°C for 20 mins. 1 μl each of Library Synthesis D Buffer and Library Synthesis D Enzyme were subsequently added, and library synthesis was performed at 22 °C for 40 mins. Immediately after, 25 μl of Library Amplification D Buffer, 1 μl of Library Amplification D Enzyme, 4 μl of nuclease-free water and 5 μl of unique Illumina-compatible indexed primer were added to each sample. Library amplification was performed using the following cycling conditions: For denaturation - 72°C for 3 mins, 85°C for 2 mins and 98°C for 2 mins, addition of unique indexes - 4 cycles of 98°C for 20 secs, 67°C for 20 secs and 72°C for 10 secs, library amplification - 14 cycles of 98°C for 20 secs and 72°C for 10 secs. Post-PCR clean-up was performed on amplified libraries with SPRI beads using 0.6X left/1x right double size selection, washed twice gently in 80% ethanol and eluted in 10–12 μl 10 mM Tris pH 8.0.

## DATA ACCESSIBILITY

A list of publicly available data used in this study can be found in **Table S1** which include data from the following studies (Cattoglio et al., 2019; Chakraborty et al., 2023; Luo et al., 2020; Thompson et al., 2022). FASTQ and processed data can be found in GEO under accession number GSE271760. Chromosome conformation capture data can also be navigated at: resgen.io/pedrorocha/FGFs/views/

## ACKNOWLEDGEMENTS

We would like to thank all members of the Unit on Genome Structure and Regulation for comments and discussions on this project and manuscript. We also thank the following people for discussions and comments on the project: Karl Pfeifer, Judy Kassis, Joana Vidigal, Nestor Saiz, Andrew Copp, Todd Macfarlan, and Tom Misteli. We also thank Anders Hansen for help establishing RCMC. We thank NICHD’s Molecular Genomics Core, specifically Fabio Faucz, Tianwei Li, and James Iben. This work utilized the computational resources of the NIH HPC Biowulf cluster (http://hpc.nih.gov). We thank the entire NICHD animal facility, and specifically Victoria Biggs for mouse husbandry, and Alex Grinberg and Jeanne Yimdjo for zygotic electroporations. For the Micro-CT analyses, we thank Louise Lanoue from the Mouse Biology Program and Douglas J. Rowland from the Center for Molecular and Genomic Imaging, both at University of California, Davis, CA, United States. This work was funded by NIH intramural project HD008975. This project has also been funded in part with Federal funds from the National Cancer Institute, National Institutes of Health, under Contract No. HHSN261201500003I. The content of this publication does not necessarily reflect the views or policies of the Department of Health and Human Services, nor does mention of trade names, commercial products, or organizations imply endorsement by the U.S. Government.

## AUTHOR CONTRIBUTIONS

SC, NW, MJA, AE, MB, JJT, PA, TJP and PPR performed experiments. PA and RC designed and generated transgenic mouse lines. SC, NW, MJA, JJT, AAE, LMSA, RKD, ML, and PR analyzed data. SC and PR wrote the manuscript with input from all the authors.

---

**Figure S1.**
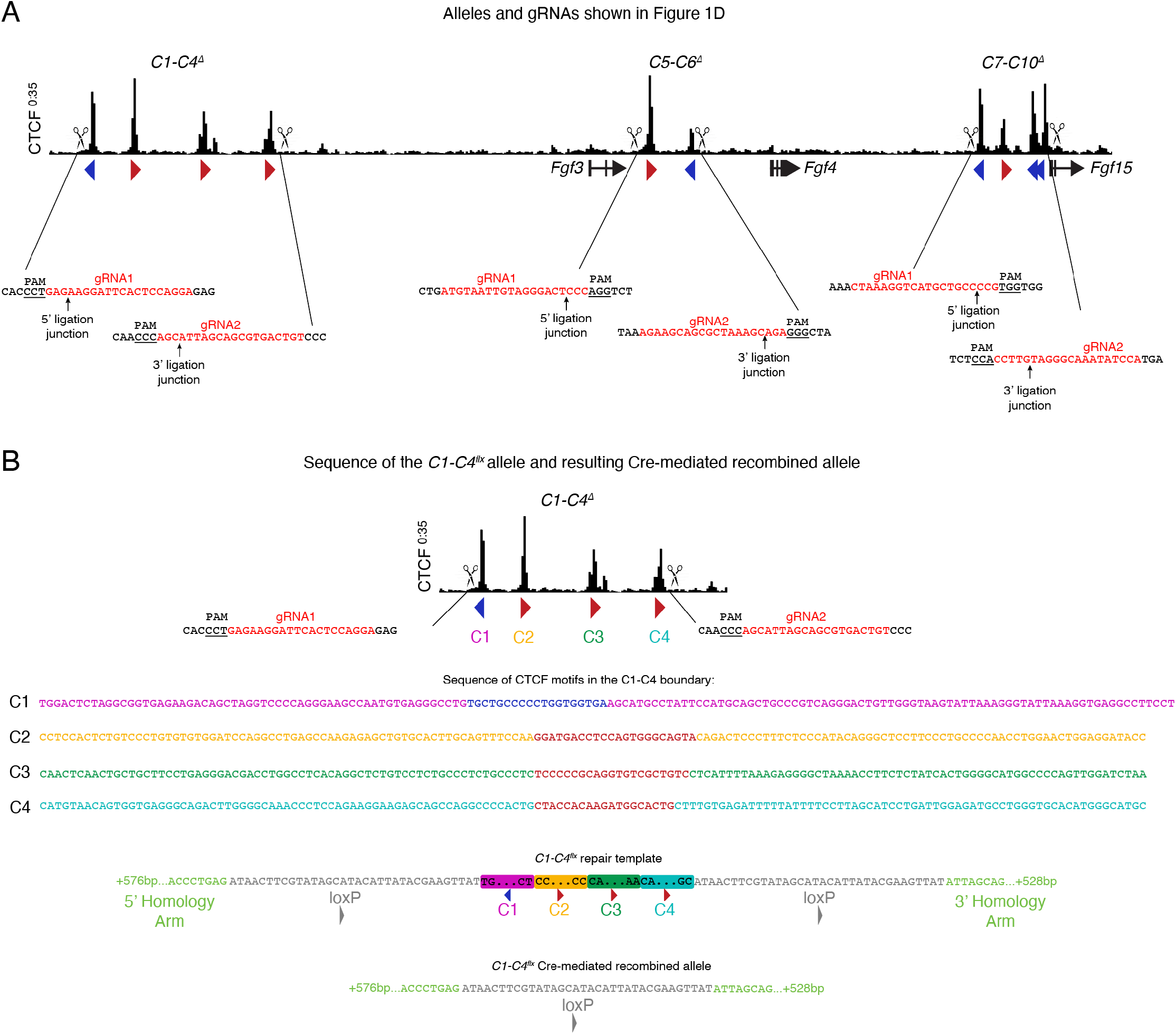
Schemes of alleles and gRNAs used to generate mouse lines in this study. **A** gRNAs used to generate the three mouse lines shown in Figure 1. PAM is shown underlined. Arrow shows location of ligation junctions between upstream and downstream cut site as determined by Sanger sequencing. **B** To generate *C7-C4*^*flx*^ the same gRNAs as in C1-C4 deletion were used. The repair template was generated by combining the CTCF core motif of the four CTCF motifs found within C1-C4, plus approximatelly 60bp on each side. The sequences of each of the four motifs are shown in different colors and the core CTCF motif is in blue or red depending on orientation (blue-negative strand, red-positive strand). Below sequences, the structure of the repair template containing those four motifs is shown, together with the sequences of *loxP* sites that surround them and a part of the homology arms. In the bottom, the sequence of the allele following Cre-mediated recombination is shown.

**Figure S2.**
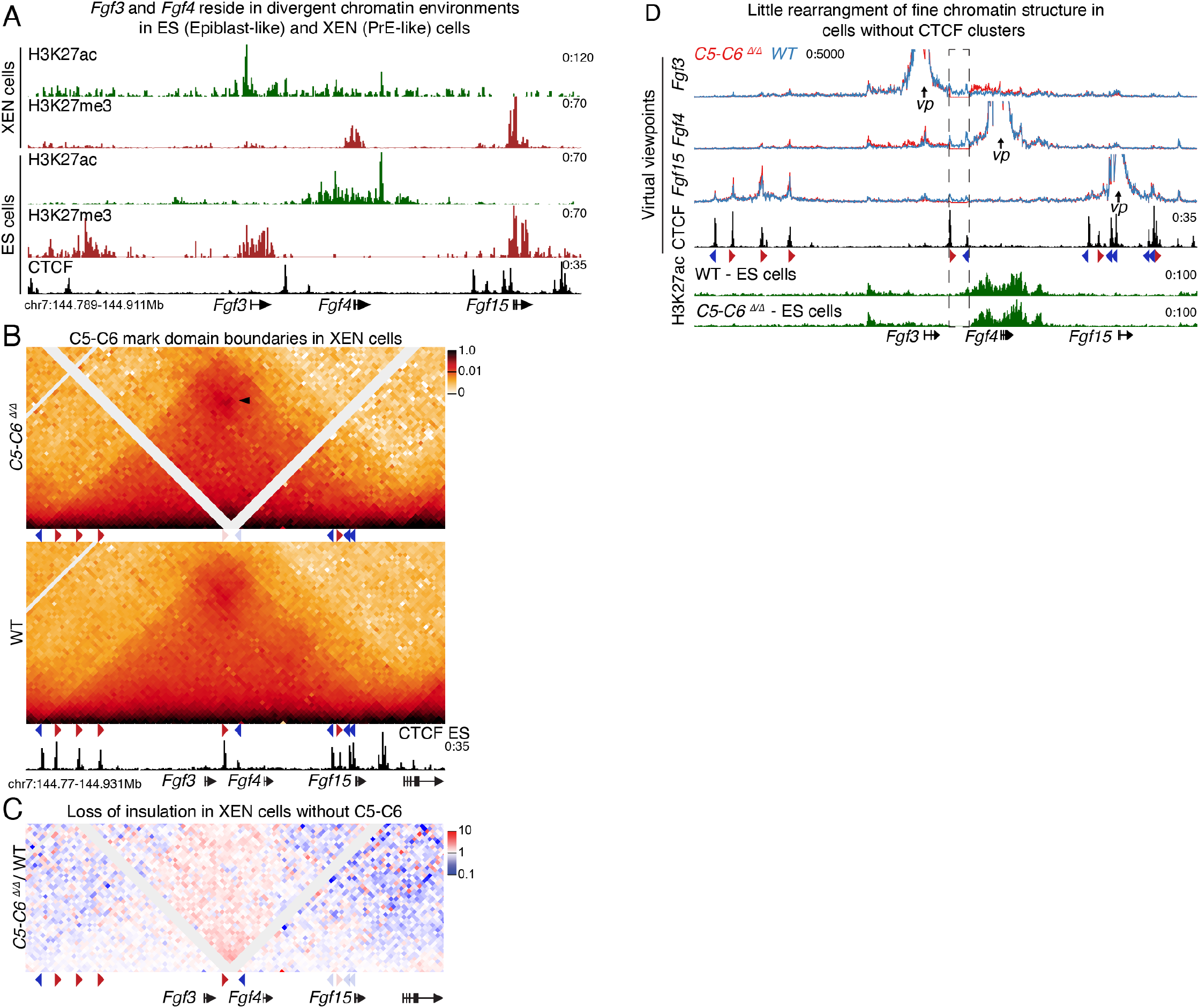
Tissue-specific expression of *Fgf3* and *Fgf4* in blastocysts does not require CTCF-mediated insulation. **A** CUT&RUN data in ES and XEN cells show that *Fgf3* and *Fgf4* have divergent patterns of H3K27ac and H3K27me3 enrichment. *Fgf4* shows active marks in ES cells and inactive in XEN cells, while *Fgf3* shows the opposite pattern **B** CHiC 1D interaction frequency heatmap in homozygotic *C5-C6*^*Δ/Δ*^ homozygous XEN cells, compared to WT at 2kb resolution. Arrow-head represent increased focal interactions between the CTCF clusters that surround the deleted cluster. **C** Differential CHiC interaction frequency heatmap. Red signal represents interactions that occur at higher frequency in mutant cell lines compared to control and blue shows interactions of lower frequency. **D** RCMC data shown as virtual viewpoints, from either *the Fgf3, Fgf4* and *Fgf15* viewpoints at 50bp resolution.

**Figure S3.**
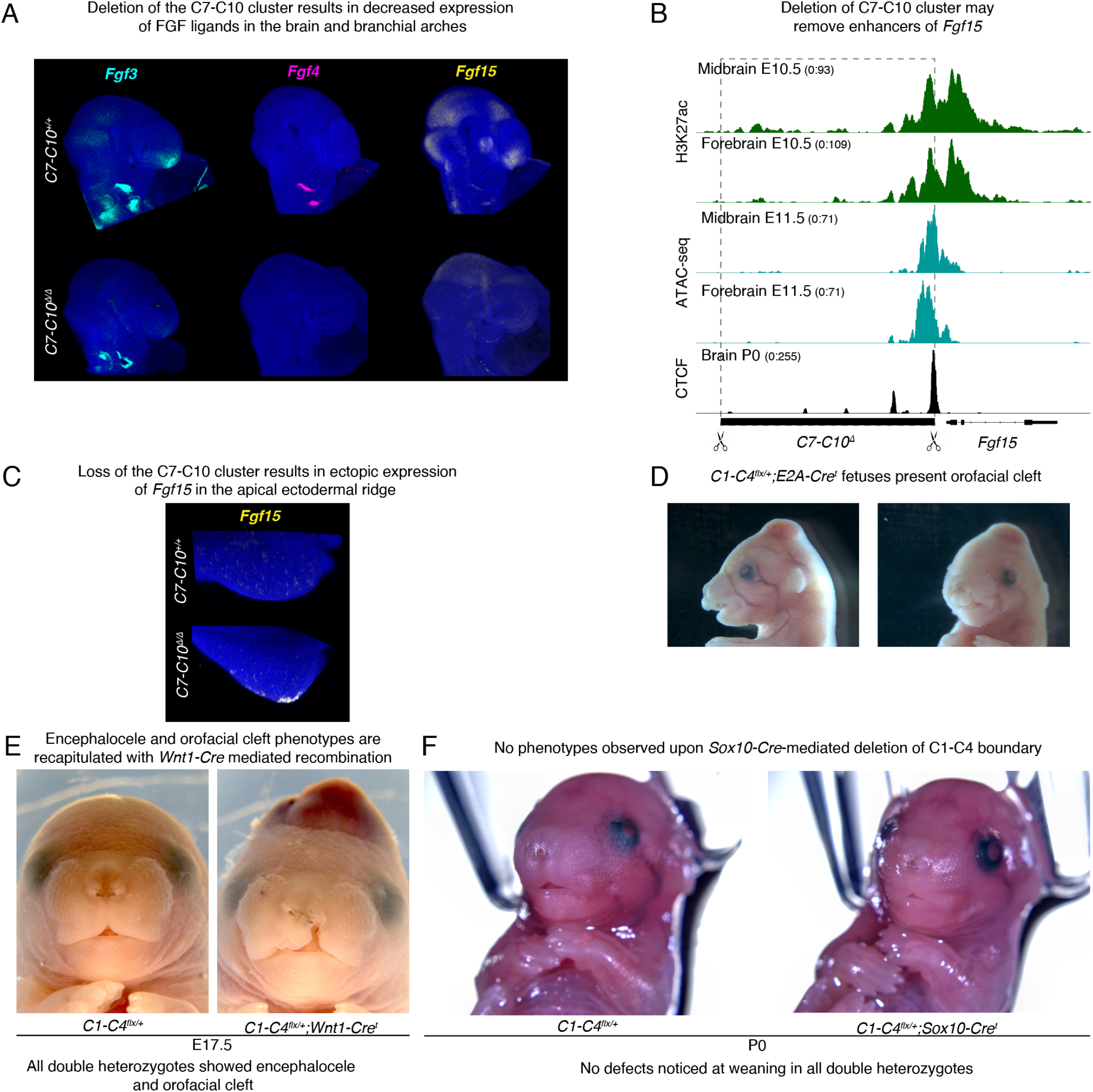
Examination of developmental phenotypes in mice without CTCF clusters surrounding the 3 FGF genes. **A** Expression of the 3 FGF genes are reduced in E9.5 embryos in *C7-C10*^*Δ/Δ*^ homozygotes analyzed by HCR (n=3/3). **B** ChlP-seq for H3K27Ac and ATAC-seq in embryonic brain shows overlapping enhancer elements and open chromatin regions with CTCF motifs are deleted in in *C7-C10*^*Δ/Δ*^ homozygotes. **C** Ectopic expression of *Fgf15* in E9.5 lorelimbs of *C7-C10*^*Δ/Δ*^ homozygotes analyzed by HCR (n=2/2 limbs).**D** Fetuses from the breeding of heterozygous *C1-C4*^*flx/+*^ with *E2A-Cre*^*t/t*^ homozygotes fully recapitulate the phenotypes seen in *C1-C4*^*Δ*^. At E16.5, these fetuses show the same orofacial cleft phenotype seen in mice with *C1-C4*^Δ^fetuses (n=6/6). E Heterozygous *C1-C4*^*flx/+*^ mice were crossed with *Wnt1-Cre*^*t*^ hemizygous mice. Double heterozygous fetuses showed orofacial cleft and encephalocele phenotypes as fetuses with germline deletion (n=6/6). **F** Heterozygous *C1-C4*^*flx*/+^ were crossed with *Sox10-Cre*^*t*^ hemizygous mice. No cleft or encephalocele were observed in progeny (10/10)

**Figure S4.**
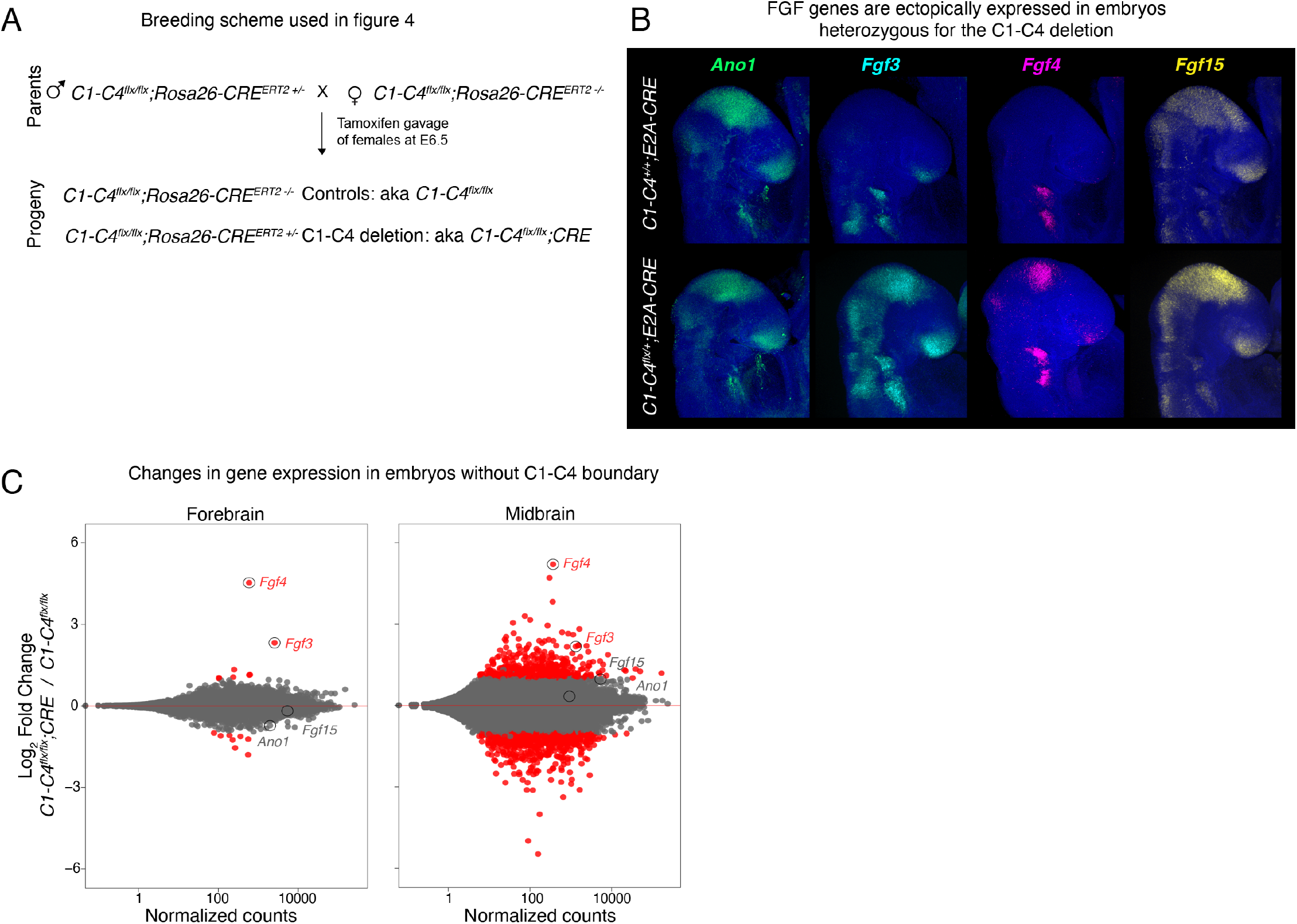
Disruption of the C1-C4 boundary leads to ectopic expression of FGF genes in the brain. **A** Scheme of the breeding and tamoxifen treatment used to generate mice shown in Figure 4. **B** HCR of E9.5 embryos showing higher expression of FGF genes in the midbrain and anterior forebrain in heterozygous embryos with C1-C4 deletion (n=3 for each genotype). **C** MA-plot of E11.5 anterior forebrain and midbrain showing all differentially expressed genes between the two genotypes. Red shows genes that were considered to be differentially expressed (adjusted pvalue<0.i, log2FC>2).

**Figure S5.**
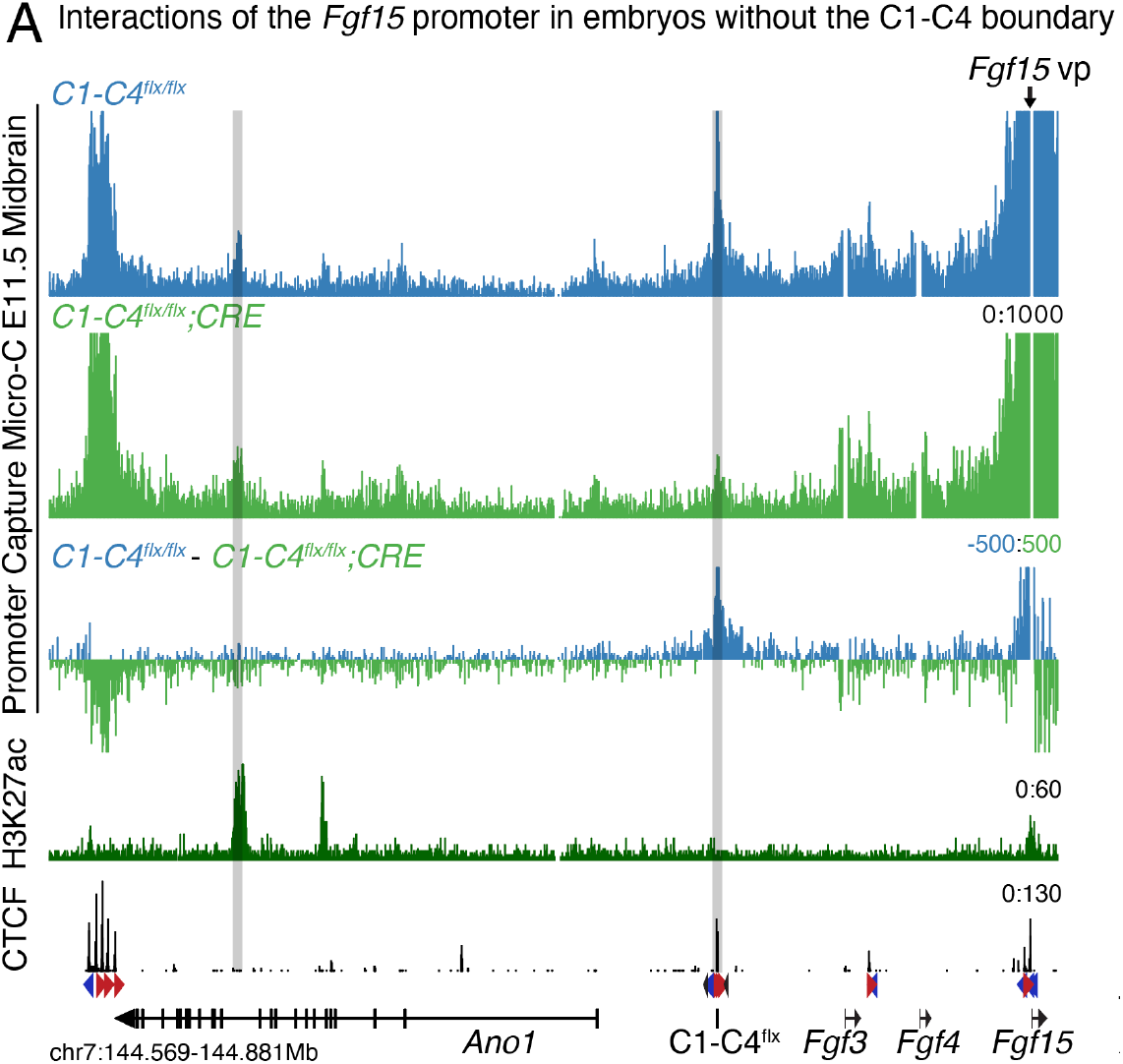
Loss of CTCF-mediated insulation exposes FGF genes to distal brain enhancers of *Ano1*. **A** Promoter Capture Micro-C shown at 50bp resolution from the *Fgf15* promoter. Gray highlight shows interactions between the promoters and the *C1-C4*^*flx*^ rescue cassette and interactions with putative *Ano1* brain enhancer.

## Notes

https://www.ncbi.nlm.nih.gov/geo/query/acc.cgi?acc=GSE271760

https://resgen.io/pedrorocha/FGFs/views

### Competing Interest Statement

The authors have declared no competing interest.

